# The transcriptomic landscape of spinal V1 interneurons reveals a role for En1 in specific elements of motor output

**DOI:** 10.1101/2024.09.18.613279

**Authors:** Alexandra J. Trevisan, Katie Han, Phillip Chapman, Anand S. Kulkarni, Jennifer M. Hinton, Cody Ramirez, Ines Klein, Graziana Gatto, Mariano I. Gabitto, Vilas Menon, Jay B. Bikoff

## Abstract

Neural circuits in the spinal cord are composed of diverse sets of interneurons that play crucial roles in shaping motor output. Despite progress in revealing the cellular architecture of the spinal cord, the extent of cell type heterogeneity within interneuron populations remains unclear. Here, we present a single-nucleus transcriptomic atlas of spinal V1 interneurons across postnatal development. We find that the core molecular taxonomy distinguishing neonatal V1 interneurons perdures into adulthood, suggesting conservation of function across development. Moreover, we identify a key role for En1, a transcription factor that marks the V1 population, in specifying one unique subset of V1^Pou6f2^ interneurons. Loss of En1 selectively disrupts the frequency of rhythmic locomotor output but does not disrupt flexion/extension limb movement. Beyond serving as a molecular resource for this neuronal population, our study highlights how deep neuronal profiling provides an entry point for functional studies of specialized cell types in motor output.

## INTRODUCTION

Neural circuits in the spinal cord are tasked with the formidable challenge of generating precise patterns of muscle contraction that underlie behavior. To achieve coordinated motor output, the spinal cord relies on networks of interneurons that integrate supraspinal commands and sensory information to shape the temporal dynamics of motor neuron activity. Accordingly, substantial effort has been devoted to identifying and classifying interneurons in the spinal motor system. Early classification schemes primarily relied on anatomical and functional characteristics, which revealed several canonical interneuron types including Group Ia interneurons that mediate reciprocal inhibition of antagonistic motor neurons, Renshaw cells responsible for the recurrent inhibition of motor neurons, and presynaptic inhibitory interneurons that gate incoming sensory input.^1–5^ With the advent of molecular and genetic approaches to neural circuit dissection, a core cellular taxonomy has emerged in which cardinal classes of ventral interneurons, termed V0 to V3, can be differentiated based on developmental provenance and gene expression.^6–8^ This knowledge, in turn, has provided insight into the functions of each neuronal population in limb movement, most prominently in the context of locomotion.^7,9–15^ Yet the extent of spinal interneuron subtype diversity, the mechanisms contributing to their diversification, and their functional roles in the motor system remain enigmatic, even as it becomes increasingly clear that each interneuron class is itself highly heterogeneous.^16,17^ Elucidating these issues is crucial to understanding how spinal circuits shape motor output.

Among ventral neuronal classes, V1 interneurons constitute a highly diverse set of inhibitory neurons whose ablation or silencing severely perturbs motor function, resulting in slowed rhythmic locomotor output and limb hyperflexion.^10,14,18,19^ Despite V1 interneurons having a central role in motor control, we understand little about the core molecular distinctions between V1 subsets and the underlying mechanisms through which V1 heterogeneity arises, nor do we understand the functional logic through which different V1 subsets might contribute to behavior. Advances in single-cell transcriptomics have greatly facilitated the ability to resolve molecular differences in cell type identity across the nervous system.^20,21^ Initial studies on the developing and adult mouse spinal cord detailed the landscape of neuronal and non-neuronal cells, collectively painting a picture in which spinal interneuron diversity emerges through the combined actions of spatial and temporal mechanisms, such that individual dorsal and ventral progenitor domains sequentially produce distinct cell types according to a temporal code of differential expression of transcription factors (TFs).^17,21–28^ Nevertheless, a full accounting of cell types within any given interneuron class remains elusive, particularly in the mature nervous system, where it has been difficult to resolve clearly defined types of ventral interneurons using single-cell transcriptomics.^25,29^

Here, building on our prior work that identified molecularly distinct subsets of V1 interneurons on the basis of differential expression of 19 TFs,^30,31^ we take advantage of single-nucleus RNA-sequencing (snRNA-seq) to perform a deep transcriptomic analysis of V1 interneurons across postnatal development and into adulthood. Targeted enrichment of V1 interneurons enabled snRNA-seq of more than 89,000 nuclei, providing unparalleled single-cell resolution of an individual spinal interneuron class and its molecular subgroups. This analysis revealed a previously unknown V1 subset, while giving insight into the repertoire of TFs and ion channels that characterize distinct V1 subsets. An analysis of V1 interneurons across postnatal development further indicates that their extensive cellular diversity documented in the neonatal spinal cord is largely recapitulated in the adult, suggesting that core V1 cell type identity is conserved from the initial establishment of neural circuit formation to the mature nervous system. Finally, to better understand the mechanisms underlying the generation of this diversity, we performed snRNA-seq on V1 interneurons isolated from mice lacking the homeodomain TF Engrailed1 (En1), which marks the V1 population.^32^ This uncovered a key role for En1 in specifying a highly restricted subset of V1 interneurons. Interestingly, while ablation of V1 interneurons results in multiple motor deficits, including slowed rhythmic locomotor output and limb hyperflexion,^10,14,18^ loss of En1 in V1 interneurons only slowed rhythmic locomotor output while preserving normal flexion/extension limb movements, thereby dissociating two core aspects of motor output. Beyond serving as a resource for studying V1 interneurons in the mouse spinal motor system, this transcriptomic dataset will enhance our understanding of neuronal subtype diversity on a broader scale, including in comparative studies with other species that may provide insight into evolutionary conservation and species-specific innovations in spinal interneuron identity.^33^

## RESULTS

### Generation and validation of a V1 transcriptomic dataset

Despite their critical role in motor output, V1 interneurons constitute only a small fraction of the overall cellular ecosystem in the spinal cord.^22^ We therefore devised a genetic strategy to selectively enrich for lineage-traced V1 interneurons, opting to perform single-nucleus rather than single-cell analysis due to several advantages: (1) Single-nucleus analysis can be performed on postnatal and adult spinal cord tissue, from which viable cells are difficult to isolate;^28^ (2) it avoids experimental artifacts from stress-induced transcriptional changes that occur within intact cells during dissociation;^34^ and (3) it is an effective means of accurately resolving cell-type identity.^35^ We used the INTACT (isolation of nuclei tagged in specific cell types) method, in which GFP^+^ V1 nuclei were isolated by fluorescence-activated cell sorting (FACS) from *En1::Cre; RC.lsl.Sun1-sfGFP* mice (hereafter called *En1^INTACT^* mice) at four postnatal stages (P0, P14, P28, and P56) (Figure 1A).^36^ This resulted in an ∼60-fold enrichment in GFP^+^ V1 nuclei relative to the pre-sorted population (Figure S1A-C).

**Figure 1.**
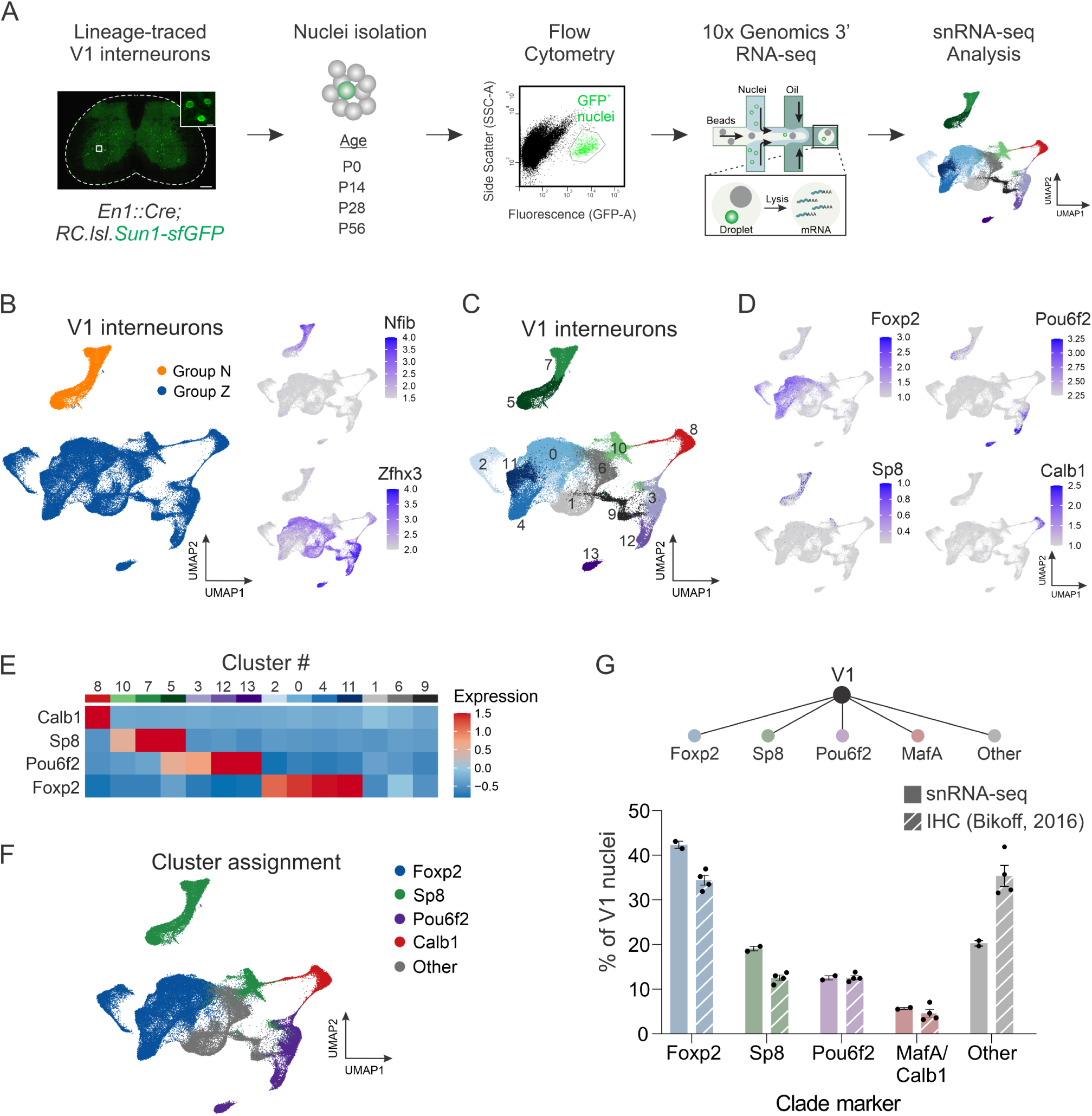
Single-nucleus transcriptomic profiling identifies V1 clades as molecularly distinct subsets. (A) Schematic of the experimental design for single-nucleus transcriptomic analysis of V1 interneurons. (B) Unsupervised clustering of V1 interneuron nuclei at low resolution reveals the first division corresponds to V1 interneurons expressing either Nfib (Group N) or Zfhx3 (Group Z). All UMAP plots depicting gene expression display log normalized values. (C) Unsupervised clustering of V1 interneuron nuclei at higher resolution identifying 14 clusters. (D) UMAP showing expression of the V1 clade markers Foxp2, Pou6f2, Sp8, and Calb1 (a proxy for MafA), which largely segregate in UMAP space. (E) Heatmap showing the scaled average expression per cluster of V1 clade markers. (F) Assignment of clade identity based on expression of Foxp2, Sp8, Pou6f2, and Calb1. (G) Comparison of the proportion of V1 interneurons contained within the clusters assigned to a given clade versus the proportion previously measured by immunohistochemical analysis of P0 lumbar spinal cord, where V1^Foxp2^ = 42.3% ± 0.8% vs 34.4% ± 1.1%, V1^Sp8^ = 19.1% ± 0.5% vs 12.5% ± 0.6%, V1^Pou6f2^ = 12.6% ± 0.5% vs 12.6% vs 0.4%, and V1^MafA/Calbindin1^ = 5.7% ± 0.2% vs 4.6% ± 0.9%; mean ± SEM; n = 2 replicates or 4 mice. All immunohistochemical data is from Bikoff et al.^30^ See also Figures S1-S4.

We next performed droplet-based snRNA-seq using the 10x Genomics platform (v3.1, see Table S1 for CellRanger quality control (QC) metrics). Although our FACS approach yielded strong enrichment of V1 nuclei, we still needed to ensure that only V1 interneurons were included in the subsequent analysis; thus, we developed a rigorous pipeline in which V1 nuclei were identified by both negative and positive selection criteria (Figure S1D; Methods). This enabled us to overcome two key challenges: (1) excluding any contaminating non-V1 nuclei that remained in the post-FACS sample, and (2) ensuring that we could identify V1 nuclei, given the absence of detectable mRNA expression of GFP and En1, a marker of V1 identity that is downregulated in the postnatal spinal cord. We first employed iterative negative selection criteria to remove contaminating nuclei corresponding to non-V1 neuronal populations, oligodendrocytes, astrocytes, and other non-neuronal populations (Figure S1E-H; Table S2). In parallel, we used a previously published embryonic whole spinal cord reference dataset to annotate postnatal nuclei. Toward this end, we collected and profiled V1 nuclei from embryonic day (E) 13.5 spinal cords of *En1^INTACT^* mice to obtain an age-matched sample for comparison to the reference scRNA-seq dataset containing clearly annotated V1 interneurons, as defined by expression of *En1*.^22^ Using the Seurat label-transfer algorithm, we calculated a V1 prediction score and identified embryonic nuclei that were assigned a putative V1 identity (Figure S1I-N). Comparison of the putative V1 nuclei in our postnatal dataset with the V1 nuclei in the embryonic data set enabled us to positively identify V1 nuclei with high confidence, even in the absence of detectable *En1* expression.

In total, we isolated 89,494 V1 nuclei that passed QC metrics across two independent replicates of four postnatal ages, with ambient RNA removed via SoupX^37^ (Figures 1 and S2; Table S3). These nuclei contained a mean of 6356 unique molecular identifiers (UMIs) and 2622 unique genes per nucleus (Figure S2B-D). To overcome batch effects arising from experimental sources of variability, we integrated the data using the Seurat anchor-based workflow, followed by dimensionality reduction and clustering to identify communities of cells.^38^ Unsupervised clustering at a low resolution and visualization in uniform manifold approximation and projection (UMAP) space revealed that the first division corresponds to V1 interneurons expressing either Nfib (Group N) or Zfhx3 (Group Z), consistent with prior observations that a basic axis of organization for ventral interneurons centers on medial, late-born, local-projecting (Group N) neurons versus lateral, earlier-born, long-distance projecting (Group Z) neurons (Figure 1B).^39^ To identify cell-type diversity at a more refined level, we subjected our V1 dataset to higher resolution clustering and identified 14 distinct clusters that ranged in size from 22.9% (Cluster #0) to 2.2% (Cluster #13) of the larger V1 population (Figures 1C and S2F).

We previously found that V1 interneurons segregate into at least four mutually exclusive subsets (clades) based on immunohistochemical analysis of the TFs Foxp2, MafA, Pou6f2, and Sp8.^30^ Accordingly, we next explored whether clades delineated on the basis of single TFs indeed represent distinct clusters when assessed by their overall transcriptomic profiles. *Foxp2*, *Pou6f2*, and *Sp8* were detected at sufficient levels to assign cluster identity based on the z-score of the average expression per cluster (Figure 1D-F), with *Foxp2* (clusters #0, #2, #4, and #11) and *Pou6f2* (clusters #3, #12, #13) robustly detected in postnatal V1 interneurons, and *Sp8* (clusters #5, #7, and #10) more weakly detected. Due to low detection of *MafA*, *Calbindin1* (*Calb1*) was used as a proxy marker for the *MafA* clade (cluster #8), which likely corresponds to Renshaw cells. Notably, these clades represented almost completely mutually exclusive clusters, with only cluster #5 showing some degree of *Pou6f2* and *Sp8* co-expression (Figure 1E), indicating that a small fraction of V1^Sp8^ and V1^Pou6f2^ neurons may be similar at the molecular level. In total, analysis of all 89,494 nuclei identified only 39 nuclei in which clade markers were co-expressed, confirming the mutually exclusive nature of these markers in postnatal V1 interneurons and their segregation into molecularly distinct V1 subsets. Moreover, the proportion of each V1 clade in our snRNA-seq data (assessed based on cluster assignment rather than per nucleus to account for the high dropout rate of snRNA-seq) was approximately comparable to those assessed via immunohistochemical analysis (Figure 1G).

In prior work that relied on TF co-expression data and spatial information, we used a sparse Bayesian framework to infer the existence of multiple cell types within each V1 clade.^31^ Therefore, we asked whether this general logic was observed in our snRNA-seq data, and used the V1^Sp8^ clade as a test case, focusing on the expression of the Sp8-associated TFs Prox1, Prdm8, and Oc2, and St18, a TF implicated in specifying projection neuron identity in the globus pallidus^40^ (Figure S3). This analysis revealed two clusters (#5 and #7) with various combinations of *Prox1*, *Prdm8*, and *St18* expression, and a single cluster (#10) expressing *Oc2* but lacking *Prox1*, *Prdm8*, and *St18*, demonstrating general alignment with predictions from the Bayesian model, albeit with fewer identified cell types (Figure S3B-F). Given the discrete nature of the V1^Sp8/Oc2^ cluster, we sought to validate its presence via immunohistochemical analysis. Although we observed few V1^Sp8^ neurons expressing *Oc2* (2.3% ± 0.4%) in mid-lumbar (L3/L4) spinal segments of P0 mice, approximately 40% of all V1^Sp8^ neurons in L5 segments were positive for *Oc2*, confirming the co-expression patterns observed in this V1^Sp8^ cluster and highlighting rostrocaudal distinctions in interneuron identity (Figure S3G).^41,42^ Together, these data show that V1 clades distinguished by *Foxp2*, *Pou6f2*, *Sp8*, and *MafA/Calbindin* expression represent molecularly distinct V1 cell types.

### Identification of a novel V1 subset

Although the four primary V1 clades accounted for nearly 80% of all V1 interneurons in the snRNA-seq dataset, a substantial portion (clusters #1, #6, and #9) falls outside of those groups. Therefore, we sought to identify unique markers that would provide genetic access to the remaining V1 interneurons. Screening of differentially expressed genes identified *Rnf220*, an E3 ubiquitin ligase implicated in the specification of spinal progenitor domains and modulation of AMPA receptor-mediated synaptic transmission, as highly selective for clusters #1 and #6 (Figure 2A).^43,44^ Immunohistochemical analysis confirmed the expression of Rnf220 in V1 interneurons (Figure 2B), with V1^Rnf220^ neurons constituting ∼13% of the overall V1 population and occupying a fairly broad spatial distribution within the ventral spinal cord (Figure 2C-D). To determine whether V1^Rnf220^ neurons comprise a non-overlapping subset with the other four clades, we performed pairwise immunohistochemical analysis of Rnf220 with Foxp2, Pou6f2, Sp8, or MafA in the presence of lineage-traced V1 interneurons (Figure 2E). Quantification revealed that V1^Rnf220^ neurons represent a largely non-overlapping population that is distinct from V1^Pou6f2^ and V1^MafA^ clades but modestly overlaps with V1^Foxp2^ and V1^Sp8^ clades (Figure 2F-G). No single marker was expressed exclusively in the sole remaining cluster (#9); however, those neurons were most specifically characterized by expression of *Nr5a2* (Figure S3A and Table S4). Thus, the snRNA-seq data leads us to posit the existence of a fifth V1 clade, marked by the expression of *Rnf220*, and indicates that more than 95% of all V1 interneurons can be assigned to one of these five clades (Figure 2H).

**Figure 2.**
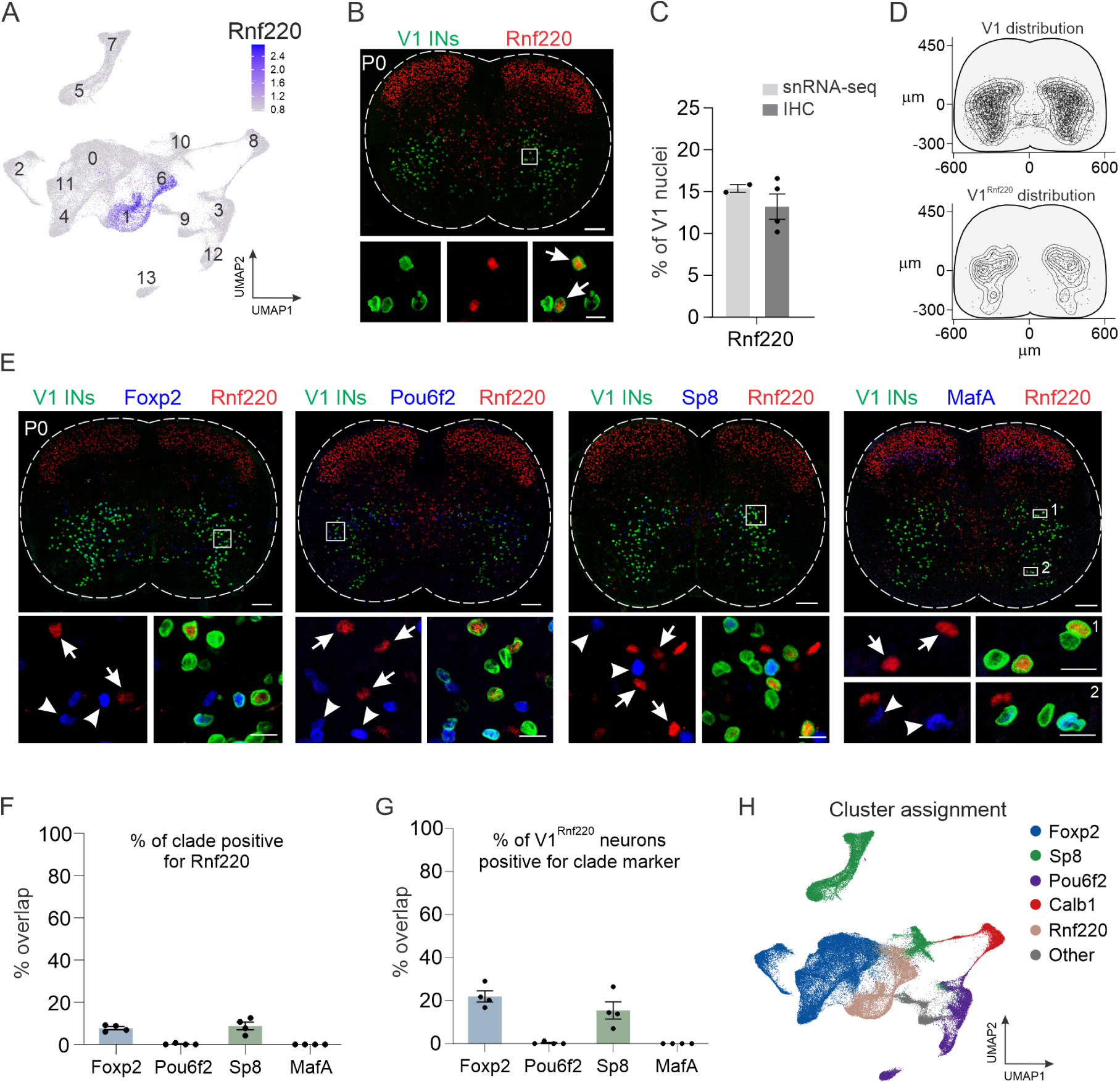
Identification of a novel V1 interneuron subset. (A) Expression of *Rnf220* within clusters #1 and #6, which are not defined by the other four clade markers. (B) Immunohistochemical analysis of Rnf220 expression in lumbar spinal cord of P0 *En1^INTACT^* mice, demonstrating expression in V1 interneurons (arrows). Scale bar = 100 μm (top) or 20 μm (bottom). (C) Fraction of V1^Rnf220^ interneurons in P0 lumbar spinal cord assessed by the proportion of V1 interneurons in Rnf220^+^ clusters #1 and #6 (14.9% ± 0.5%) or via immunohistochemistry (13.2% ± 1.5%); mean ± SEM; n = 2 snRNA-seq replicates or 4 mice, respectively. (D) Spatial distribution of V1^Rnf220^ interneurons largely recapitulates the distribution of the overall V1 population. (E-G) Rnf220-expressing V1 interneurons represent a largely non-overlapping population with the four major clades, completely distinct from V1^Pou6f2^ and V1^MafA^ clades and exhibiting modest overlap with V1^Foxp2^ and V1^Sp8^ clades. Scale bars = 100 μm (top) or 20 μm (bottom) (H) Assignment of V1^Rnf220^ clusters relative to V1^Foxp2^, V1^Sp8^, V1^Pou6f2^, and V1^Calb1^ clusters, which together constitute 95% of all V1 nuclei.

### Molecular profiles of V1 interneuron clades

Having established that five V1 clades can be distinguished by their transcriptomic profiles, we next sought to explore their overall taxonomy and molecular differences. Our prior classification schema based on limited TF expression implied that cell types within a clade are more similar than cell types across clades. Yet the opposite may be true, particularly when cells are assessed by their overall transcriptomic profile. Hierarchical clustering analysis revealed that certain clusters within a clade are more transcriptomically similar to each other than to clusters from other clades (e.g. clusters #5 and #7 within the V1^Sp8^ clade, or clusters #4 and #11 within the V1^Foxp2^ clade) (Figure S4A). This rule, however, was not absolute. For example, cluster #0 in the V1^Foxp2^ clade was most similar transcriptomically to cluster #10 in the V1^Sp8^ clade. Moreover, we identified highly distinct molecular profiles within a single clade, best exemplified by the V1^Pou6f2^ clade, in which clusters #12 and #3 were very similar, but cluster #13 was located at the far end of the dendrogram. The number of differentially expressed genes (DEGs) per cluster (Figure S4B and Table S4), as well as Pearson correlation and distance in principal component space (Figure S4C-D), confirmed that cluster #13 is highly distinct. Thus, V1 subsets within an individual clade are often but not always closely aligned on the molecular level.

To better understand the unique molecular signatures of V1 clusters, we next focused on two broad categories of genes, TFs and ion channels, due to their important roles in shaping cellular identity and physiological properties. Analysis of the most differentially expressed TFs showed robust cluster-specific cohorts of TF genes, including many of the 19 TFs previously identified and validated as subdividing the larger V1 population^30^ (Figures 3A and S3A). For example, cluster #8, which corresponds to Renshaw cells, was defined not only by coincident expression of *Oc1*, *Oc2*, *MafA*, and *MafB* (Figure S3A), which is consistent with previous evidence,^45^ but also by expression of *Tle4*, *Nfia*, *Rarb*, and *Ppargc1a*, all of which have been implicated in aspects of nervous system development.^46–49^ In addition to *Prox1*, *Prdm8*, *Oc2*, and *St18*, the V1^Sp8^ clade was characterized by enriched expression of several TF families: Nfi-family members *Nfib* and *Nfix*, which control cellular differentiation in neurons and glia;^50,51^ Ebf-family members *Ebf1* and *Ebf3*, which couple neuronal differentiation to cell-cycle exit in the spinal cord;^52^ and Tshz-family members *Tshz2* and *Tshz3*, the latter of which is implicated in cortico-striatal synaptic transmission and autism-like behavior.^53^ The V1^Pou6f2^ clade, previously shown to express Nr5a2, Lmo3, MafB, Oc1, and Oc2, was characterized by several additional TFs involved in neuronal specification: Pbx-family members *Pbx1* and *Pbx3*, which function as Hox cofactors and mediate segregation and clustering of spinal motor neurons within motor columns;^54^ *Zfp462*, a Pbx1-interacting TF that safeguards neuronal lineage specification via epigenetic regulation of heterochromatin;^55,56^ *Pou2f2*, which regulates the spatial organization of spinal V2a interneurons; *Maml3*, a regulator of Notch signaling,^57^ and *Zfhx3*, which is associated with long-range projection neurons in the spinal cord.^39^ The largest V1 clade, V1^Foxp2^ interneurons, previously shown to express Foxp1/2/4, Otp, Nr4a2, Esrrb (Nr3b2), Esrrg (Nr3b3), and Lmo3 in varying combinations, was also enriched for *Pax2*^58^, *Casz1*, *Nr3c2*, and *Dach1*. Finally, V1^Rnf220^ interneurons were characterized by enriched expression of *Pax8*, previously suggested to participate in dorsal inhibitory interneuron specification,^59^ Bcl11-family members *Bcl11a* and *Bcl11b* (cluster #6), and *Esrrg*, *Nr3c2*, and *Dach1* (cluster #1). RNAScope analysis of select TFs, including *Pax8*, *Bcl11a*, *Dach1*, and *St18*, independently validated their expression in V1 interneurons (Figure 3B). Together with earlier immunohistochemical validation of many of these differentially expressed V1 TFs,^30^ this analysis provides insight into the transcriptional programs differentiating subsets of V1 interneurons.

**Figure 3.**
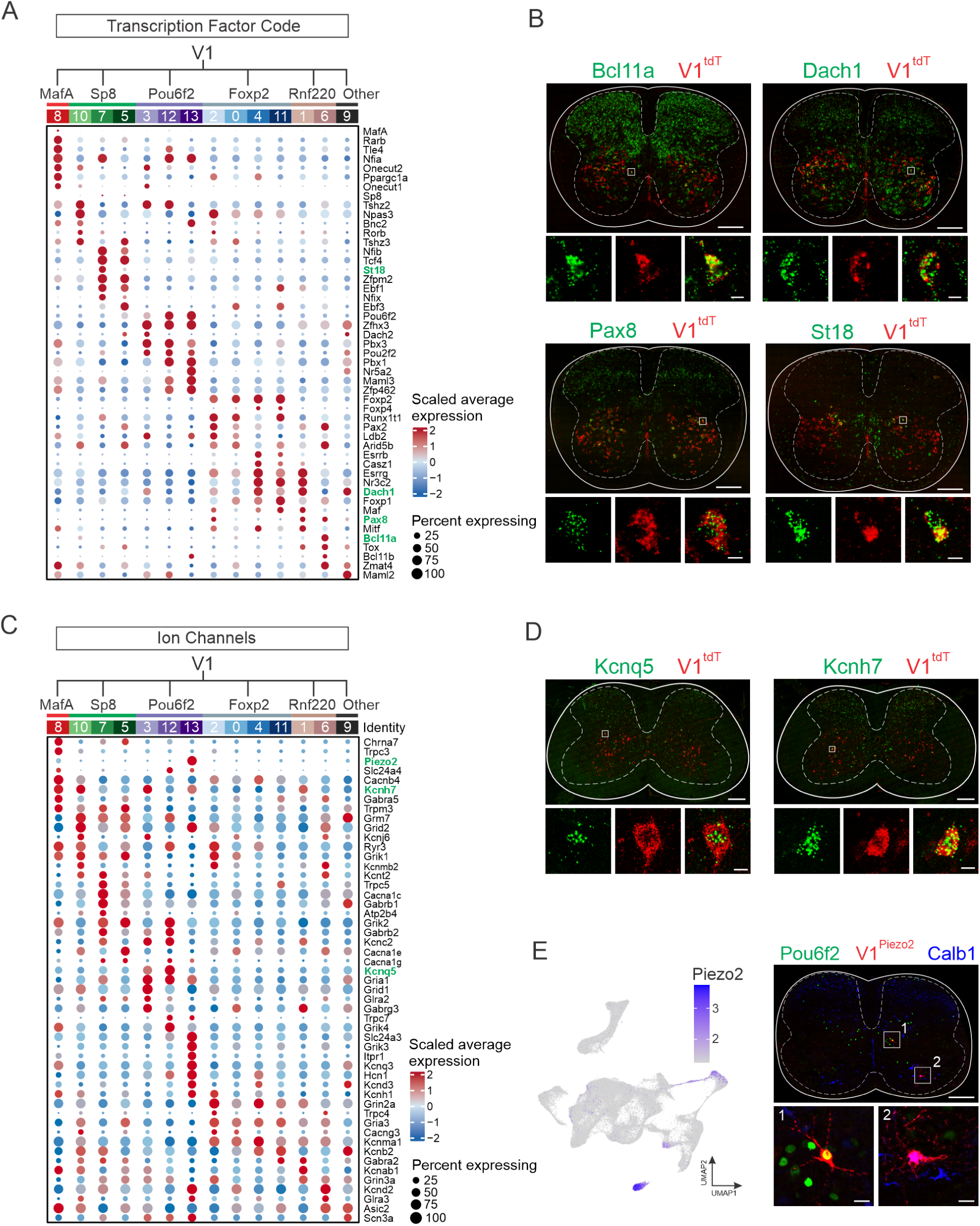
Molecular profiles of V1 interneuron subsets. (A) Average scaled expression of transcription factors (TFs) enriched in each of the 14 V1 clusters. TFs validated by *in situ* hybridization (ISH) appear in green text. (B) RNAScope ISH validating expression of select TFs in V1 interneurons in P0 lumbar spinal cords of *En1::Cre; Ai14* mice. Scale bars = 200 μm (top) or 10 μm (bottom). (C) Average scaled expression of ion channels enriched in each of the 14 V1 clusters. Validated genes appear in green text. (D) RNAScope ISH validating expression of select ion channels in V1 interneurons in P28 lumbar spinal cord of *En1::Cre; Ai14* mice. Scale bars = 200 μm (top) or 10 μm (bottom). (E) Left, UMAP showing *Piezo2* expression in V1 interneurons. Right, P0 cervical spinal cord from *Piezo2::Cre; En1::Flpo; Ai65D* mouse demonstrating Pou6f2 expression (green) or Calb1 expression (blue) in V1^Piezo2^ interneurons (red). Of the V1^Piezo2^ neurons in cervical segments, 41.1% ± 11% were Pou6f2^+^, 36.1% ± 4.5% were Calb1^+^, and 22.8% ± 6.6% were not positive for either marker, whereas at lumbar segments 2.8% ± 1.9% were Pou6f2^+^, 58.0% ± 2.1% were Calb1^+^, and 39.2% ± 2.0% were not positive for either marker (mean ± SEM, n = 3 animals). Scale bars = 200 μm (top) or 20 μm (bottom). See also Figures S3 and S4.

We next examined the differential expression of ion channels and transporters amongst subgroups of V1 interneurons, which have distinct electrophysiological signatures.^30^ Members of the voltage-gated K^+^ channel family were particularly enriched, with 13 of the top DEGs belonging to this family (Figure 3C). Ionotropic glutamate receptors (10 of the top DEGs), Trp channels (5 of the top DEGs) and voltage-gated Ca^2+^ channels (5 of the top DEGs) also constituted a substantial portion of identified ion channels. RNAScope *in situ* hybridization (ISH) of select differentially expressed KCN-family members that regulate neuronal excitability (e.g. *Kcnq5*^60^ and *Kcnh7*^61^) confirmed the expression of these genes in V1 interneurons (Figure 3D). Interestingly, we found that *Piezo2*, the principal mechanoreceptor mediating light touch and proprioception,^62,63^ was among the most highly selectively expressed ion channels, with particularly strong expression in cluster #13, a subset of the V1^Pou6f2^ clade, and weak expression in cluster 8, representing the MafA/Calb1 clade (Figure 3C,E). To determine if *Piezo2* is indeed expressed in subsets of V1 interneurons, we generated *Piezo2::Cre; En1::Flpo; Ai65D* mice to lineage-trace V1^Piezo2^ interneurons. We identified tdTomato-expressing neurons that were positive for *Pou6f2* expression and *Calb1* expression in cervical spinal cords of P0 mice, suggesting that these neurons are mechanosensitive (Figure 3E). Collectively, our snRNA-seq data reveals that in addition to their unique TF profiles, V1 clades express distinct cohorts of ion channels that may in part reflect differences in their electrophysiological properties.

### Changes in gene expression across postnatal development do not alter the core identity of V1 interneuron clades

Following their birth, spinal neurons undergo a protracted period of maturation that extends from embryogenesis well into adulthood.^64^ Embryonic V1 interneurons segregate into distinct cell types shortly after neurogenesis,^22^ consistent with observations that V1 clades are born in sequential but partly overlapping waves from E9.5 to E13.5^45,65,66^. By P0, they exhibit even more extensive molecular diversity.^31^ Yet it remains unclear how this developmental diversity relates to mature neural circuits in the spinal cord. One possibility is that the diversity of neonatal V1 interneurons represents a transient stage in the development of motor circuits, analogous to how some *Drosophila* olfactory projection neurons exhibit peak diversity during circuit assembly but then converge to largely indistinguishable cell types in adults.^67^ This possibility is suggested by the observation that V1 interneurons (and ventral interneurons more generally) in the adult spinal cord exhibit less pronounced distinctions in cell type identity than do their dorsal interneuron counterparts.^25,28,29^ Alternately, such diversity may be maintained in mature circuits, analogous to interneurons in the adult cortex.^68^

To explore whether V1 interneurons retain their transcriptomic identity, we assessed the extent to which V1 interneurons from different postnatal ages contribute to each of the clusters. V1 interneurons from P0, P14, P28, and P56 mice were found in each cluster, in roughly equal proportions (Figure 4A-B), indicating that new V1 subtypes do not appear with age, nor does the initial diversity present at birth decrease with age. Consistent with this finding, quantification of the contribution of V1 nuclei to each cluster, as a function of age by using integrated local inverse Simpson’s index (iLISI), showed a high degree of intermixing (Figure 4C). Together, these data suggest that the core identity of V1 interneurons in neonatal mice is preserved through adulthood.

Despite the overall consistency in V1 cell-type identity, we observed substantial age-related changes in gene expression (Figure 4D and Table S5). The largest changes occurred during the first 2 postnatal weeks, a period characterized by rapid maturation of synaptic connectivity, myelination, and the development of intrinsic cell properties.^69^ In contrast, changes were less pronounced from P14 onward. Gene ontology (GO) analysis of the most differentially expressed genes at P0 identified enrichment of those involved in synapse organization and neuron projection morphogenesis/development, whereas at P28 and P56 DEGs were associated with synaptic transmission and transsynaptic signaling (Figure 4E). Consistent with these cellular processes, we identified decreasing developmental expression of *Sema6d*, which is involved in guiding axons to their appropriate targets^70^, and increasing expression of *Snap25*, a key component of synaptic vesicle exocytosis machinery (Figure 4F).^71^ Beyond global changes in V1 interneuron transcriptomic profiles across development, we identified cluster-specific changes in gene expression, as exemplified in cluster #8, where *Piezo2* expression was highest at P0 and downregulated by P14 onward, whereas in cluster #13, *Piezo2* expression had the opposite dynamic, showing increased expression from P0 to P56 (Figure 4G). Thus, V1 interneurons exhibit clear developmental changes in gene expression during postnatal maturation yet maintain stability of their general cell-type identity.

**Figure 4:**
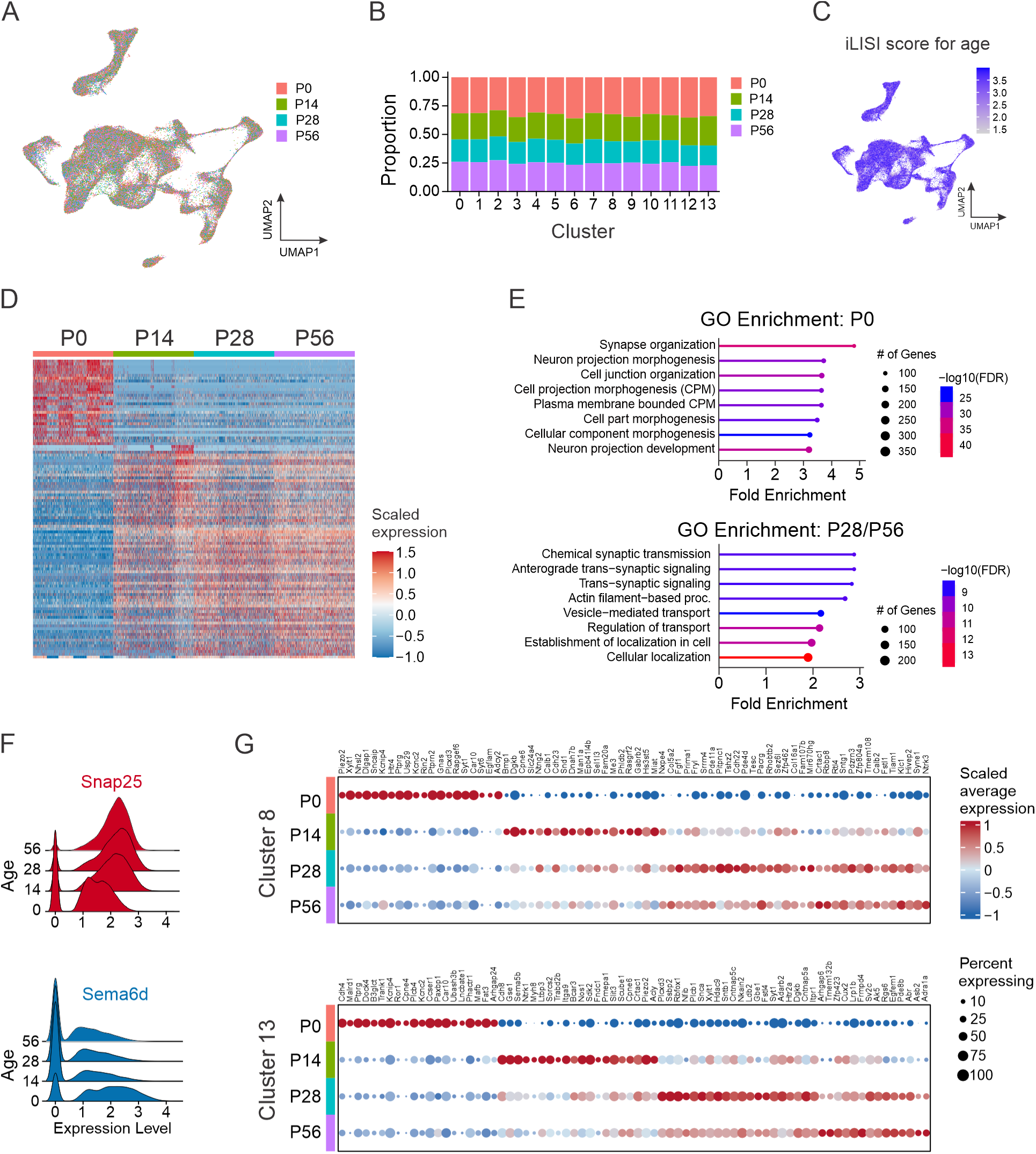
Changes in gene expression across postnatal development do not change core neuronal identity. (A) UMAP of V1 nuclei color-coded by age (P0, P14, P28, or P56) at the time of sample collection. (B) The relative contribution of each sample to the 14 V1 clusters. (C) Integrated local inverse Simpson’s index (iLISI) calculated for age. iLISI scores ranged from 1 to 4 (the number of groups), with higher scores indicating better intermixing between groups. (D) Heatmap of the top 30 most differentially expressed genes at each time point. (E) Gene ontology (GO) analysis showing the top 8 biological processes enriched in the gene expression profiles of V1 nuclei at P0 (top) or P28 and P56 combined (bottom). (F) Ridge plot highlighting genes with increasing expression (*Snap25*, red) and decreasing expression (*Sema6d*, blue) from P0 to P56. Log normalized values are shown. (G) Identification of genes dynamically regulated across postnatal development that are specific to cluster #8 (top) or Cluster #13 (bottom).

### Transcriptomic analysis of V1 interneurons in *En1* KO mice

We next considered whether the TF En1 itself controls the development of V1 interneurons. This question was motivated by prior observations that homozygous *En1^taulacZ^* mice lacking En1 protein exhibit a dramatic slowing of rhythmic locomotor-like output,^10^ yet they do not have obvious deficits in V1 interneuron fate specification.^72,73^ To explore En1 function, we took advantage of the fact that the *En1::Cre* allele disrupts the endogenous *En1* locus,^74^ thereby creating a null allele and generating *En1* knockout (*En1^KO^*) animals in the homozygous state (Figure 5A, top). Consistent with prior observations, the overall number of V1 interneurons was comparable in *En1^Het^* and *En1^KO^* animals (268 ± 9 neurons and 265 ± 7 neurons per 20 μm section, respectively; mean ± SEM, n = 5 mice, *p* = 0.815, unpaired two-tailed t-test) (Figure S5A). Following isolation of V1 nuclei from the spinal cords of P0 *En1^Het^* and *En1^KO^* mice, we performed snRNA-seq and isolated 7,553 nuclei that passed QC, revealing 11 distinct clusters (Figure 5A, bottom). The molecular identity of these clusters aligned well with our earlier postnatal V1 dataset (Table S6), though we identified fewer clusters which may reflect limitations in resolving more subtle gene expression differences, given the 10-fold lower sampling of nuclei. Segregation of the data by genotype revealed one cluster (#10) with a paucity of *En1^KO^* nuclei (Figure 5B). Quantification revealed a striking reduction (∼2.5-fold decrease) of *En1^KO^* nuclei in cluster #10 in two independent biological replicates (Figure 5C). In contrast, the proportion of *En1*^Het^:*En1^KO^* nuclei within each of the other 10 clusters was nearly equivalent, indicating the selective paucity of neurons in cluster #10 (Figure 5D). Moreover, our analysis of TF expression in *En1^Het^* and *En1^KO^* mice showed no substantial differences in gene expression, indicating cell-type identity of the remaining V1 interneurons in *En1^KO^* mice is not significantly perturbed (Figure S5B).

**Figure 5.**
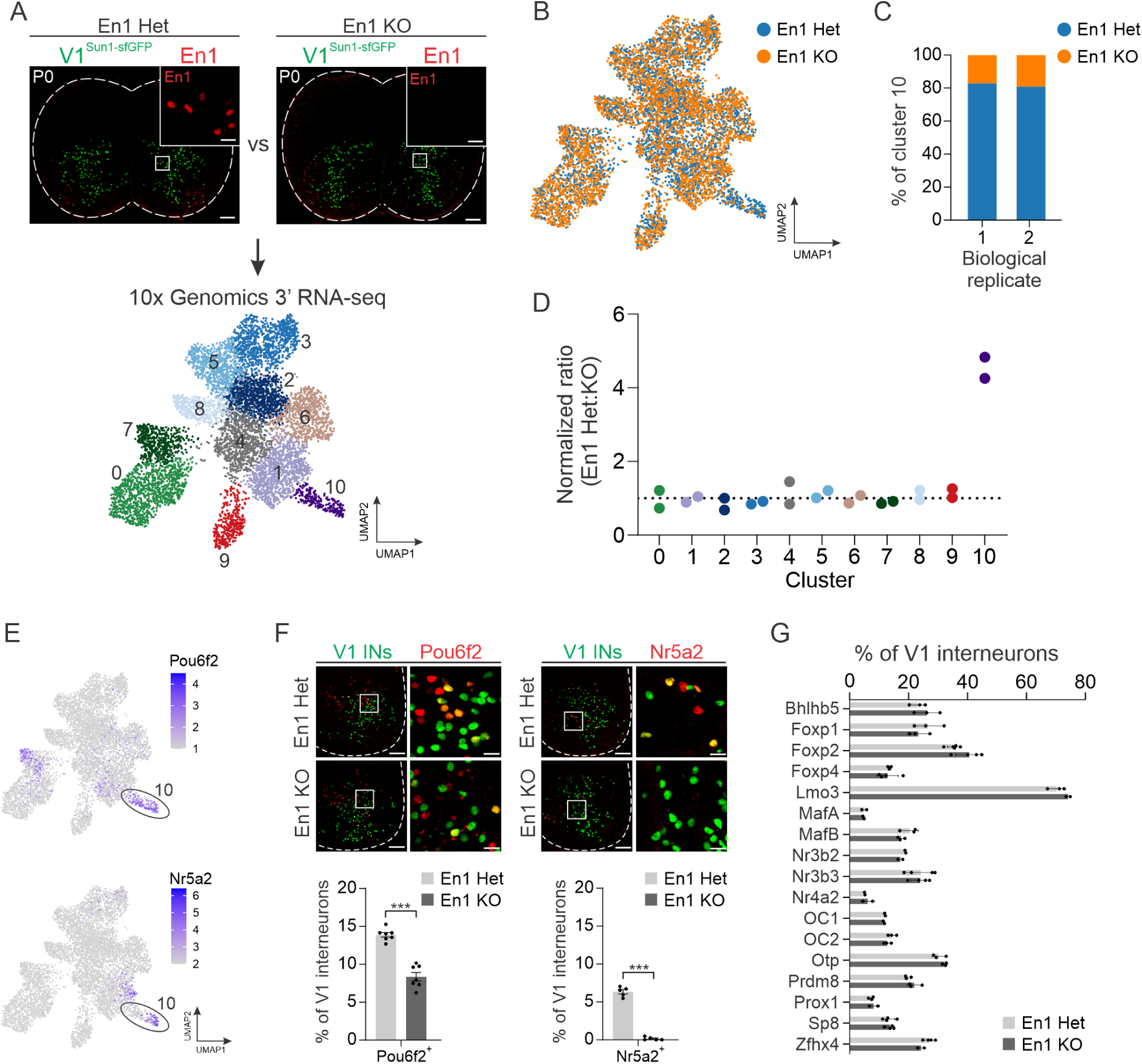
Transcriptomic analysis of V1 interneurons in *En1^KO^* mice reveals En1 is required for the development of a specific V1^Pou6f2^ cell type. (A) Top, representative images of P0 lumbar spinal cords from *En1::Cre* heterozygous; *RC.lsl.Sun1-sfGFP* (En1 Het) or *En1::Cre* homozygous; *RC.lsl.Sun1-sfGFP (*En1 KO) mice immunostained for En1 (red) and lineage-traced V1 interneurons (green). Bottom, UMAP visualization of unsupervised clustering of snRNA-seq data generated for these genotypes, identifying 11 distinct clusters. (B) UMAP of *En1^Het^* (blue) and *En1^KO^* (orange) nuclei. (C) Normalized percentage of nuclei in cluster 10 originating from either *En1^Het^* or *En1^KO^* animals across two biological replicates. (D) Normalized ratio of Het:KO nuclei in each cluster across two biological replicates, where 1 corresponds to equal representation of both genotypes. With cluster #2 as a reference, only cluster #10 was found to have a credible change in abundance, as determined by scCODA (log2FC = −1.61, False Discovery Rate [FDR] < 0.05). (E) UMAP showing expression of *Pou6f2* and *Nr5a2* enriched in Cluster #10. (F) Immunohistochemical (IHC) analysis of V1^Pou6f2^ (left) and V1^Nr5a2^ (right) interneurons in P0 lumbar spinal cords of *En1^Het^* or *En1^KO^* mice demonstrating a significant reduction in each subset. V1 interneurons were lineage-traced using *Tau.lsl.nLacZ* mice (green). V1^Pou6f2^: 13.9% ± 0.3% vs 8.4% ± 0.6% in Het vs KO animals, respectively, mean ± SEM, n = 7 animals; V1^Nr5a2^: 6.3% ± 0.3% vs 0.1% ± 0.1% in Het vs KO animals, mean ± SEM, n = 5 animals, *** *p* < 0.0001, unpaired two-tailed T-test). (G) Percentage of V1 subsets defined by 17 other TFs in *En1^Het^* or *En1^KO^* mice assessed by IHC, showing no statistical difference between genotypes (mean ± SEM, n = 2 to 5 animals per condition, *p* > 0.5 for all comparisons, unpaired two-tailed *t*-test with Holm-Sidak correction for multiple comparisons). See also Figure S5.

An analysis of V1 clade markers revealed that cluster #10 corresponded to a subset of the V1^Pou6f2^ clade that also expresses Nr5a2, suggesting there may be a substantially smaller proportion of these neurons in *En1^KO^* mice (Figure 5E). Independent immunohistochemical analysis of P0 lumbar spinal cord confirmed a significant reduction in the percentage of Pou6f2^+^ and Nr5a2^+^ V1 interneurons in *En1^KO^* mice (Figure 5F). Moreover, pseudo-bulk analysis of gene expression comparing *En1^Het^* and *En1^KO^* nuclei identified *Nr5a2* and *Pou6f2* as the two most significantly down-regulated genes in *En1^KO^* nuclei (Figure S5C). In comparison, immunohistochemical analysis of the proportion of V1 subsets defined by 17 other TFs showed no significant difference between *En1^Het^* and *En1^KO^* animals (Figure 5G). Consistent with these observations, analysis of the spatial location of V1 interneurons in lumbar spinal segments of control mice and *En1^KO^* mice revealed that the position of V1^Pou6f2^ neurons was selectively perturbed in the absence of *En1*, whereas the distribution of other V1 populations was unaffected (Figure S5D). Thus, En1 plays a critical role in the development of a specific cluster of neurons within the V1^Pou6f2^ clade.

### Functional analysis of En1 knockout mice dissociates deficits in locomotor frequency from limb hyperflexion

Given the highly selective loss of a small subset of V1 interneurons in *En1^KO^* animals, we next considered the functional implications of the lack of spinal En1 expression on motor output. More specifically, we considered whether the loss of En1 fully accounts for the previously described functional phenotypes observed upon ablation or silencing of the V1 population, namely a significantly reduced frequency of rhythmic locomotor output^10,19^ and severe hyperflexion of limbs,^14,18^ or whether these phenotypes could be dissociated from one another. We first confirmed that acute loss of V1 interneurons severely perturbs both locomotor frequency and limb flexion. As expected, ablation of V1 interneurons via administration of diphtheria toxin (DT) to *En1::Flpo; Hoxb8::Cre; Ai65D; Tau::dsDTR* mice significantly slowed locomotor rhythm, as assessed via ventral root recordings during fictive locomotion (Figures 6A-B, S6A-C). V1 ablated animals also showed severe hyperflexion of their hindlimbs, which were assessed using DeepLabCut (DLC) to quantify ankle and knee angle during the tail suspension assay (Figure 6C-D and Video S1). In contrast, the forelimbs in V1 ablated animals did not exhibit hyperflexion (Figure S6D), consistent with the incomplete recombination efficiency of *Hoxb8::Cre* in more rostral (cervical) spinal segments.^75^

**Figure 6.**
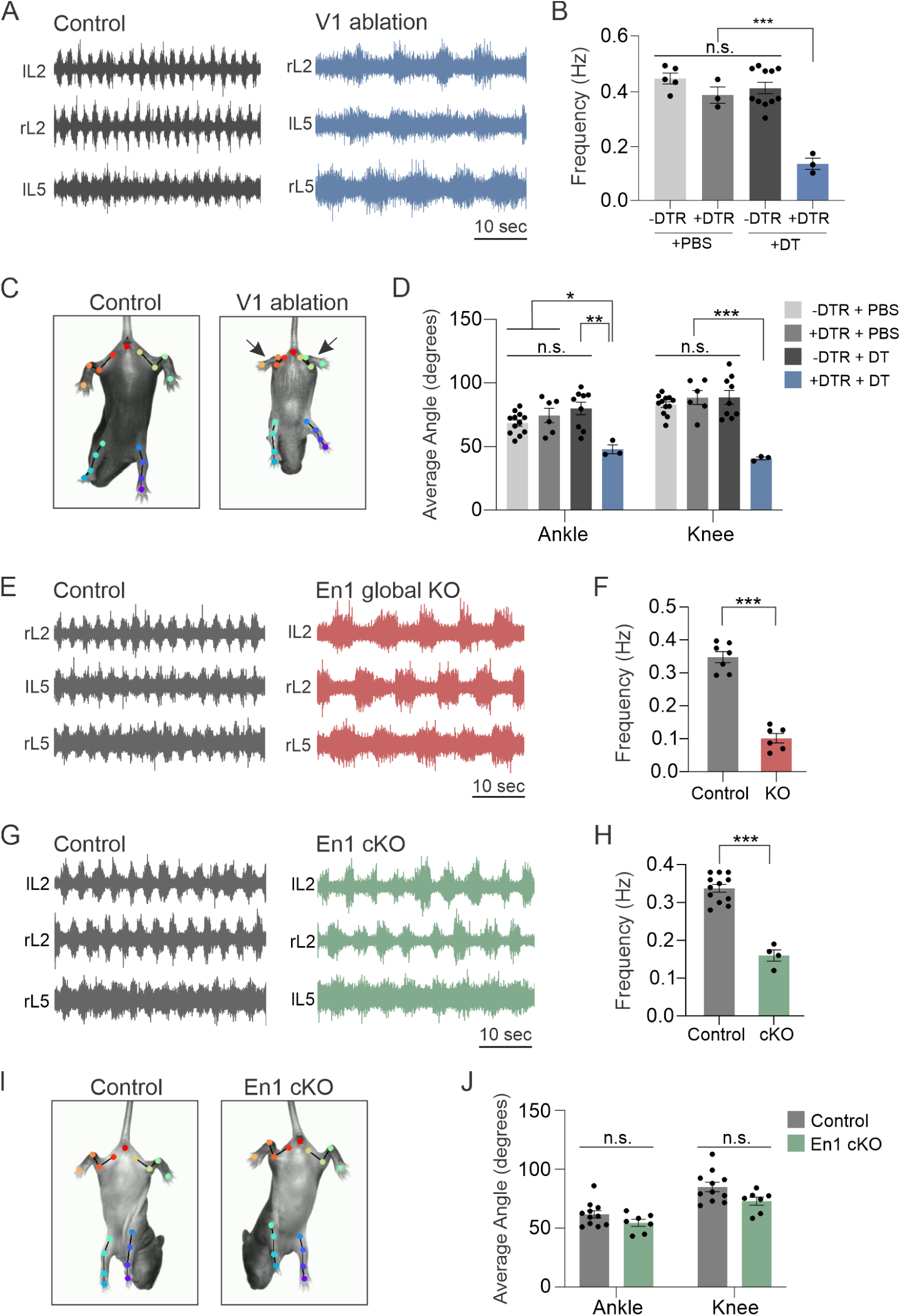
Loss of spinal En1 expression reduces the frequency of fictive locomotion but does not result in limb hyperflexion. (A) Ventral root recordings during fictive locomotion performed on P4 control (-DTR + DT, gray) or V1-ablated (+DTR + DT, blue) spinal cords. (B) Reduced frequency of fictive locomotor activity upon DT-mediated ablation of V1 interneurons (locomotor frequency of 0.44 ± 0.02 Hz, 0.38 ± 0.03 Hz, 0.41 ± 0.02 Hz in control conditions vs 0.13 ± 0.02 Hz upon V1 ablation, mean ± SEM, n = 3-11 mice, two-way ANOVA: F(1,18) = 13.86, \*\*\**p* < 0.001, two-way ANOVA followed by Holm-Sidak test for multiple comparisons). (C) Tail suspension assay performed in P7 control (-DTR +DT, left) or V1 ablated (+DTR +DT, right) mice. Hindlimb hyperflexion is indicated by arrows, with joints and angles identified using DeepLabCut. (D) The average angles of the ankle and knee joints are significantly reduced upon ablation of V1 interneurons, indicating a hyperflexed state (n = 3 to 12 mice, mean ± SEM, \**p* < 0.05, \*\**p* < 0.01, \*\*\**p* < 0.001, two-way ANOVA followed by Holm-Sidak test for multiple comparisons). (E) Ventral root recordings during fictive locomotion in control (*En1::Cre^Het^*) or global *En1* KO (*En1::Cre^Hom^*) mice. (F) Reduced frequency of fictive locomotor activity in global *En1* KO mice (n = 6-7 mice, mean ± SEM, \*\*\**p* < 0.001, unpaired two-tailed *t*-test). (G) Ventral root recordings in control (*En1^flox/+^*, gray) or *En1* cKO (*Hoxb8::Cre; En1^flox/flox^*, green) mice. (H) *En1* cKO mice show reduced locomotor frequency (n = 4-10 mice, mean ± SEM, \*\*\**p* < 0.001, unpaired two-tailed *t*-test). (I) The hindlimbs of control (*En1^flox/+^*) or En1 cKO mice observed during tail suspension assay, with normal flexion/extension. (J) The average angles of the ankle and knee joints are not significantly different upon loss of En1 (n = 7-11 mice, mean ± SEM, n.s. = not significant by unpaired two-tailed *t*-test). See also Figure S6.

To directly confirm that the loss of En1 similarly affects the speed of rhythmic locomotion, we recorded ventral root activity during fictive locomotion and observed a dramatic reduction in frequency in *En1^KO^* mice relative to *En1^Het^* animals (Figure 6E-F). However, due to neonatal lethality observed in *En1^KO^* mice, which lack En1 expression globally, we were unable to properly assess limb flexion in these animals. To bypass this neonatal lethality, we generated *Hoxb8::Cre; En1^flox/flox^* conditional knockout mice (hereafter called *En1 cKO* mice) in which deletion of En1 was restricted to the spinal cord. Immunohistochemical analysis confirmed that *En1 cKO* animals showed a complete loss of En1 protein in lumbar spinal segments (Figure S6F-H). Similar to V1-ablated and *En1^KO^* animals, *En1 cKO* animals showed a reduced frequency of locomotor-like activity duing fictive locomotion (Figure 6G-H). Importantly, however, *En1 cKO* animals exhibited normal hindlimb position during the tail suspension assay, showing none of the severe hyperflexion characteristics of V1-ablated animals (Figure 6I and Video S1). DLC tracking and analysis of limb kinematics confirmed that the mean ankle and knee angles were comparable between control and *En1 cKO* animals (Figure 6J). Together, these data indicate that loss of En1 causes a subset of the phenotypes observed upon ablation of V1 interneurons, thereby dissociating deficits in locomotor speed from perturbations in flexion/extension movement.

## DISCUSSION

Single-cell transcriptomic approaches have greatly facilitated the systematic characterization of cell type diversity across the nervous system.^21,76^ In the mouse spinal cord, numerous transcriptomic studies have characterized the larger cellular ecosystem of neuronal and non-neuronal populations in the developing and mature spinal cord.^21–25,28,39^ These global surveys have been complemented by targeted studies of spinocerebellar neurons,^77^ V2a interneurons,^41^ and perhaps most prominently, motor neurons.^78–80^ Yet the diversity within individual classes of spinal interneurons has remained obscure, limiting our ability to dissect the synaptic and circuit architecture of spinal circuits and their contributions to behavior. Here, focusing on the cardinal class of spinal V1 interneurons, we present a comprehensive single-nucleus transcriptomic atlas of more than 89,000 nuclei, spanning development from birth to adulthood, providing insight into the molecular features that distinguish major subsets of V1 interneurons and the transcriptional dynamics of postnatal V1 interneuron maturation. Moreover, we identified a role for En1 in the development of one specific subset of V1 interneurons and show how loss of *En1* selectively perturbs locomotor frequency while sparing limb flexion. These studies illustrate how deep neuronal profiling of spinal interneurons can begin to bridge the gap between neuronal diversity and motor function.

### V1 interneuron diversity in context

Given the fundamental role of spinal circuits in sensory perception and motor output, considerable effort has gone into finding an overarching logic to the classification of spinal neurons. From a developmental perspective, neuronal diversity in the spinal cord emerges through the sequential generation of neurons from distinct embryonic progenitor domains arrayed along the dorsoventral and anteroposterior spinal axes.^26^ These three principles - genetic provenance, spatial organization, and temporal specification - have provided a complementary framework to classical physiological studies to gain a better understanding of the diversity of neurons in the spinal cord. Layered onto these classification schemes, recent transcriptomic studies have revealed that dorsal versus ventral identity contributes the most variability to gene expression,^39^ and these two general categories fundamentally differ - dorsal interneurons exhibit clear transcriptomic distinctions, and ventral interneurons show more overlapping profiles.^25,29,81^

Our deep transcriptomic analysis of a single ventral interneuron class enables an exploration of the organizing principles of interneuron diversity in relation to previous studies on V1 interneurons as well as the larger cellular ecosystem in the spinal cord. We previously found that V1 interneurons comprise a highly diverse set organized into at least four mutually exclusive clades, V1^Foxp2^, V1^MafA^, V1^Pou6f2^, and V1^Sp8^, that are collectively composed of dozens of putative cell types that can be distinguished based on TF expression, cell body position, and intrinsic physiological properties.^30,31^ Strikingly, this clade organization appears to be largely conserved across 300 million years of evolution, where V1 interneurons in *Xenopus* express Foxp2, MafA, Pou6f2, and Sp8 TFs in a manner that mirrors the mutually exclusive expression observed in mice.^33^ Here we show that these four established clades and the newly identified V1^Rnf220^ clade represent distinct cell types based on unbiased transcriptomic clustering. The fractionation of V1 interneurons into 14 distinct clusters represents a resolution that identifies robust differences in gene expression and shows excellent alignment with our previous analysis of V1 clade diversity, albeit with substantially fewer total numbers of predicted cell types.^31^ Interestingly, while some clusters within a given V1 clade were quite similar, others within that same clade were highly transcriptomically divergent (e.g. V1^Pou6f2^ cluster #13 vs. clusters #3 and #12). Given that transcriptomic distinctions in cell type identity correspond with other phenotypic modalities, including anatomical and physiological properties,^82,83^ it is reasonable to expect that individual V1 clades contain interneurons with disparate connectivity profiles and roles in motor output.

Beyond this, several core insights into the likely phenotypic characteristics of V1 interneurons emerge from our transcriptomic analysis. Previous studies identified a major axis of gene expression in ventral interneurons that was predictive of mediolateral position, birthdate, and projection patterns, where Group-N (NeuroD2 and Nfib-expressing) neurons are medial, late-born, and locally projecting, and Group-Z (Zfhx3, Zfhx4, and Foxp2-expressing) neurons are lateral, early-born, and long-distance projecting.^39^ We find that at a low level of clustering, V1 interneurons divide into Group-N neurons represented by a subset of the V1^Sp8^ clade, and Group-Z neurons comprising all other clades. This general logic aligns well with prior positional analysis of V1 interneurons^30,31,39^ and birth-dating studies showing that V1 clades emerge sequentially in overlapping waves between E10 and E12.5, with V1^MafA/Renshaw^ and V1^Pou6f2^ neurons born early, followed by V1^Foxp2^ neurons, and ending with V1^Sp8^ neurons.^66^ It is somewhat less clear how the logic of Group-N and Group-Z projection patterns applies to V1 interneurons, as both V1^Foxp2^ and V1^MafA/Renshaw^ neurons provide dense, local innervation of motor neurons, despite their Group-Z (long-distance projecting) designation.^66^

Our transcriptomic analysis of V1 interneurons from P0 to P56 also provides insight into how cell-type identity changes during postnatal maturation. The largest transcriptional changes occur during the first 2 postnatal weeks, which correspond to a rapid maturation in motor behavior that culminates in adult-like locomotion by the third week.^84^ Thereafter, even as neural circuit refinement continues, the molecular profiles of V1 interneurons stabilize. This is reminiscent of motor neuron gene expression programs that are highly dynamic from embryogenesis through P21, at which point transcriptional and chromatin-accessibility landscapes stabilize.^64^ Importantly, our integrated dataset revealed a near equal contribution of all sample ages to each cluster, indicating that the extensive cellular diversity documented within V1 interneurons in the neonatal spinal cord is largely preserved through adulthood; V1 interneuron clusters retained their core identity, while undergoing temporal changes in gene expression. Our ability to resolve differences in adult V1 interneurons may in part reflect the large number of V1 nuclei sampled in our dataset, analogous to findings that defined extensive diversity within adult motor neurons following targeted enrichment of that neuronal population.^79^ Notably, several transcriptomic studies have found that cardinal classes of ventral interneuron populations are not easily distinguishable, such that V1, V2b, and other inhibitory cardinal classes merge together.^25,29^ It is possible that even as V1 interneurons retain substantial transcriptional diversity, there is a convergence of gene expression patterns for cell types in different interneuron classes that exhibit similar anatomical or functional characteristics. Indeed, functionally and anatomically defined types of spinal interneurons may span multiple cardinal populations, exemplified by Group Ia reciprocal interneurons that are found in both the V1 and V2b populations, or propriospinal neurons that originate from dI3, V1, V2, and V3 populations.^14,85^

### Involvement of class-defining transcription factors in cell-type specification

During development, spatially and transcriptionally distinct progenitor domains oriented along the dorsoventral spinal axis give rise to cardinal classes of interneurons whose postmitotic expression of many key TFs defines their identity.^30,86^ Many of these TFs play a crucial role in cell-type specification of their respective interneuron classes. Thus, *Evx1* mutant mice exhibit abnormal V0_V_ interneuron development in which V0_V_ interneurons adopt a V1-like fate,^87^ and *Chx10* mutants show substantially reduced numbers of V2a interneurons.^88^ In other cases, cell fate specification appears largely preserved. For example, *Sim1* mutant mice produce normal numbers of V3 interneurons but exhibit abnormal patterns of migration and axon pathfinding.^89^ Prior studies on *En1* mutant mice similarly found that V1 cell fate specification is preserved, with neurons exhibiting axon pathfinding deficits and reduced Renshaw cell innervation of motor neurons.^73,90^

Here, we demonstrate that *En1* is required for the normal development of a highly specific subset of V1 neurons, defined by coincident expression of *Pou6f2* and *Nr5a2*. The observation that affected neurons constitute only a tiny fraction (∼3%) of the overall V1 population likely explains why prior studies did not identify changes in V1 cell fate. Indeed, we found that the total number of V1 interneurons is unchanged in *En1^KO^* mice. This is different from the role *En1* plays in midbrain dopaminergic neurons, where *En1* and the related family member *En2* are essential for dopaminergic neuron survival.^91^ We also did not observe significant increases in other V1 clusters in *En1^KO^* animals, though this might be expected even if V1^Pou6f2/Nr5a2^ neurons adopted an altered cell fate, given their sparsity within the V1 population. While future studies will be necessary to determine the mechanisms through which En1 controls V1^Pou6f2/Nr5a2^ identity, our findings illustrate the power of using unbiased transcriptomic approaches to gain insight into the development of rare cell types in the nervous system.

### V1 interneuron diversity and the control of movement

A hallmark of motor output is its highly adaptable nature, enabling animals to tailor the speed, strength, and trajectory of limb movements to a constantly changing environment. Insight into how spinal motor circuits achieve such flexibility has emerged from studies linking broad populations of molecularly defined neurons to distinct features of motor output, including the regulation of locomotor speed, flexion/extension of joints, and inter-limb coordination.^15^ Emerging evidence indicates that such circuits are organized in a modular nature, with different subsets of interneurons and motor neurons implementing distinct components of movement.^92–95^ Yet understanding how the vast diversity of interneurons in the mammalian spinal cord relates to discrete aspects of motor output remains a major challenge. Here, we show that in the absence of *En1*, mice exhibit a pronounced slowing of rhythmic locomotor output while retaining normal flexion/extension limb movement. This is in stark contrast to what is observed upon ablation or silencing of V1 interneurons, which results in both slowed rhythmic locomotor output and severe hyperflexion of limbs.^10,14,18,19^ Although developmental deficits in axon fasciculation or synaptic connectivity may contribute to *En1*^KO^ motor deficits,^73,90^ our finding that deficits in locomotor frequency and limb flexion can be dissociated, combined with our transcriptomic characterization of *En1 cKO* mice showing normal specification of all V1 interneurons except a highly specific subset of the V1^Pou6f2^ clade, suggest a role for V1^Pou6f2/Nr5a2^ neurons in locomotor speed. In this scenario, the control of limb flexion would primarily be mediated by a different V1 subset, most likely the V1^Foxp2^ clade, which contains neurons with the classic anatomical features of Ia reciprocal inhibitory interneurons^66^ and appears unaffected by loss of *En1*. We note that we have not yet been able to selectively target V1^Pou6f2/Nr5a2^ interneurons to directly demonstrate their involvement in the control of locomotor speed. Future studies will be required to define the relative contributions of V1^Pou6f2/Nr5a2^ neurons and altered V1 connectivity to shaping locomotor speed, as well as to clarify the contribution of other V1 subsets described in our transcriptomic dataset to the control of movement.

## ACKNOWLEDGEMENTS

We are grateful to Alfonso Lavado (St. Jude Histology and Tissue Processing Core) for help with RNAScope ISH, Kim Lowe and Jim Houston (DNB Flow Cytometry Core) for technical assistance with FACS, Peter Sims, Chaolin Zhang, and Daniel Moakley for advice during pilot single-cell transcriptomic experiments, Guoqiang Gu for sharing the rat anti-St18 antibody, and Mary Patton for advice regarding ventral root recording. We also thank Lora Sweeney, David Vijatovic, Paco Alvarez, and Suresh Jetti for discussions and comments on the manuscript. Single-nucleus RNA-sequencing was performed at the St. Jude Hartwell Center for Biotechnology, which is supported, in part, by the National Cancer Institute grant P30 CA021765. This work was also supported by the National Institute of Health grant R01NS123116 (J.B.B.), the German Research Foundation grant CRC1451 - 431549029-Z02 (G.G.), and the American Lebanese Syrian Associated Charities (ALSAC) (J.B.B.). The content of this paper is solely the responsibility of the authors and does not necessarily represent the official views of the National Institutes of Health.

## AUTHOR CONTRIBUTIONS

A.J.T. and J.B.B. conceived the project. A.J.T., K.H., and P.C. conducted transcriptomic experiments. A.J.T. analyzed the transcriptomic data, with critical guidance from V.M., M.I.G., C.R., and A.K. A.J.T. and J.H. conducted imaging and behavioral experiments. I.K. and G.G. analyzed the behavioral data. A.J.T. and J.B.B. wrote the manuscript, with input from all authors. J.B.B. obtained funding and supervised all aspects of the project.

---

**Figure S1.**
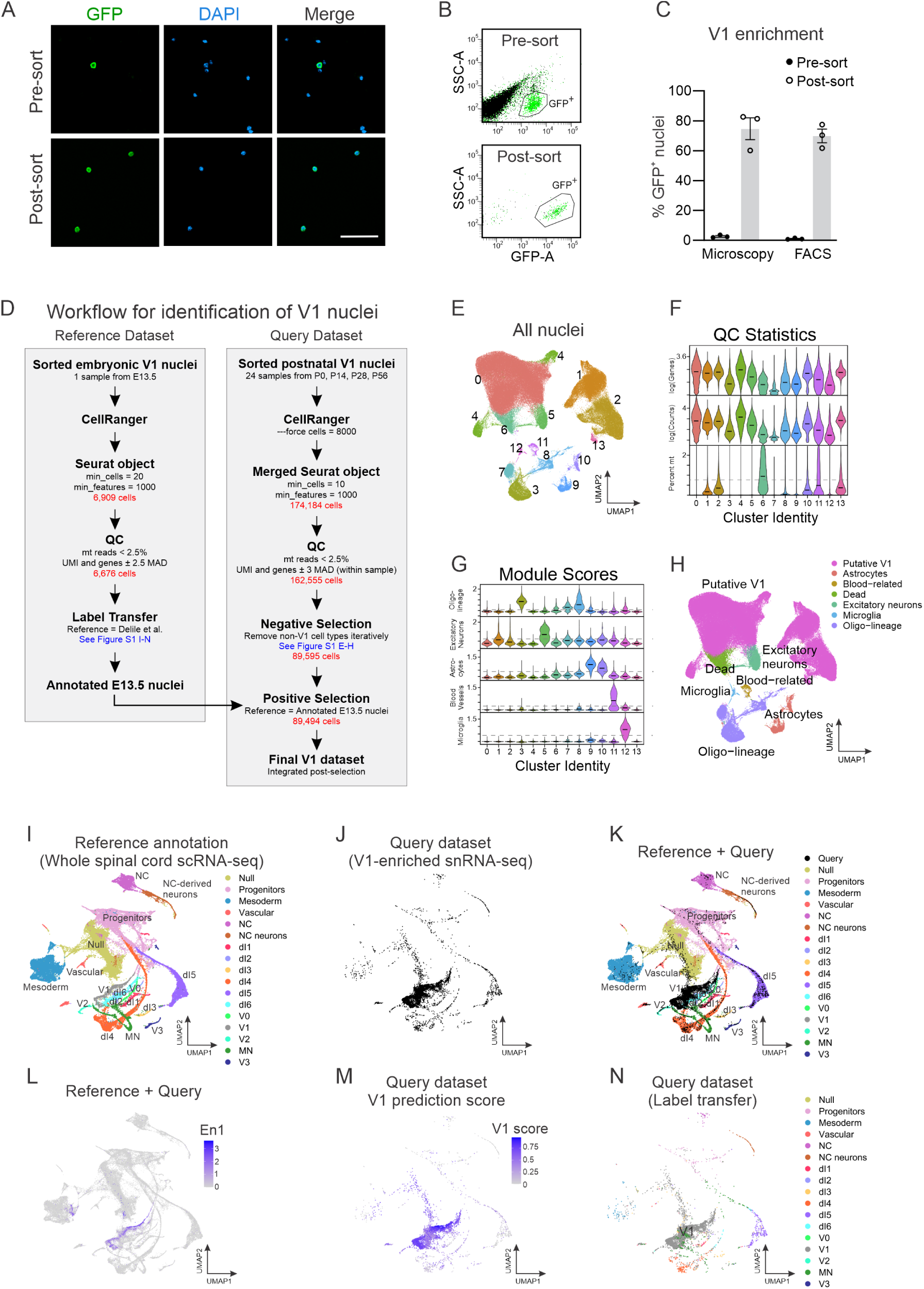
Enrichment of V1 nuclei via fluorescence activated cell sorting, and subsequent identification of V1 nuclei, related to Figure 1. (A-C) Enrichment of V1 nuclei by fluorescence activated cell sorting (FACS). (A) Example images of nuclei isolated from P0 *En1::Cre; RC::lsl.Sun1-sfGFP* (*En1^INTACT^*) spinal cords before (top) and after (bottom) FACS. Scale bar = 50 μm. B) Representative FACS analysis of spinal cord nuclei before sorting (top) and then re-analyzed after sorting (bottom). (C) Quantification of V1 nuclei enrichment assessed via microscopy or FACS before sorting (2.7% ± 0.5% and 1.1% ± 0.3%, respectively) and after sorting (74.7% ± 7.3% and 69.9% ± 4.6%, respectively) (n = 3 independent trials, mean ± SEM). (D) Workflow outlining the identification of V1 nuclei within the postnatal nuclei dataset (Query Dataset) by using both negative and positive selection criteria. (E-H) Negative selection to remove contaminating (non-V1 interneuron) nuclei. (E) UMAP of all nuclei meeting quality control (QC) cutoffs across all ages, replicates, and anatomical regions combined. Colors indicate unbiased Louvain graph-based clustering. (F) The number of genes detected per nuclei, total reads per nuclei, and percent of mitochondrial reads are shown per cluster. Cluster 6 was identified as “Dead” due to the low number of genes and counts, and high number of mitochondrial reads. (G) Gene module scores per cluster for non-V1 cell types. Clusters above the threshold (dotted line) were identified as the corresponding cell type. See Supplementary Table 1 for genes used to create the module score. (H) The UMAP shown in (E) with clusters assigned an identity per the selection measures described in (F) and (G). (I-N) Positive identification of V1 interneurons using label transfer. (I) E9.5-E13.5 whole spinal cord scRNA-seq data^22^ was used as a reference data set. (J) E13.5 *En1::Cre^INTACT^* snRNA-seq query data (black) enriched for V1 interneurons by flow cytometry. (K) Data set from (J) projected onto reference data (I). (L) Expression of the transcription factor En1 overlaid on the projected data set. (M-N) V1 label transfer prediction scores and final assignments for the E13.5 snRNA-seq data set, used for subsequent positive and negative selection of our postnatal data.

**Figure S2.**
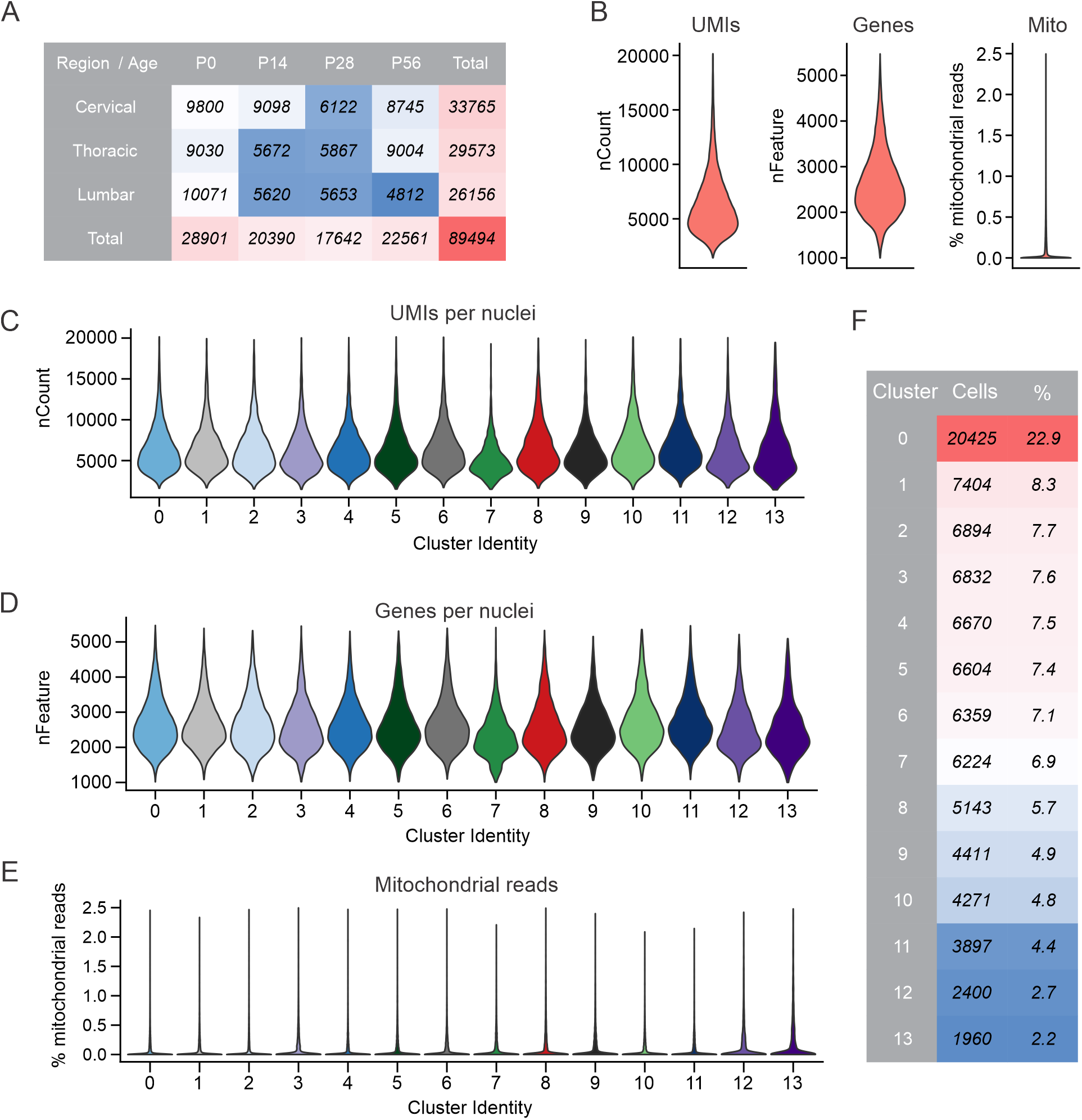
Quality Control (QC) statistics on V1 interneuron nuclei, related to Figure 1. (A) Table showing the total number of nuclei identified as V1 interneurons grouped by age and anatomical location. Colors represent low (blue) to high (red) proportions across all categories. (B) Overall QC statistics on V1 interneurons showing total number of unique molecular identifier (UMI)-corrected counts per nuclei, number of genes detected per nuclei, and the percentage of mitochondrial reads per nuclei. (C-E) Similar to (B) except the same statistics were parsed by the cluster identification shown in Figure 1C. (F) Table displaying the number and percentage of nuclei in each cluster.

**Figure S3.**
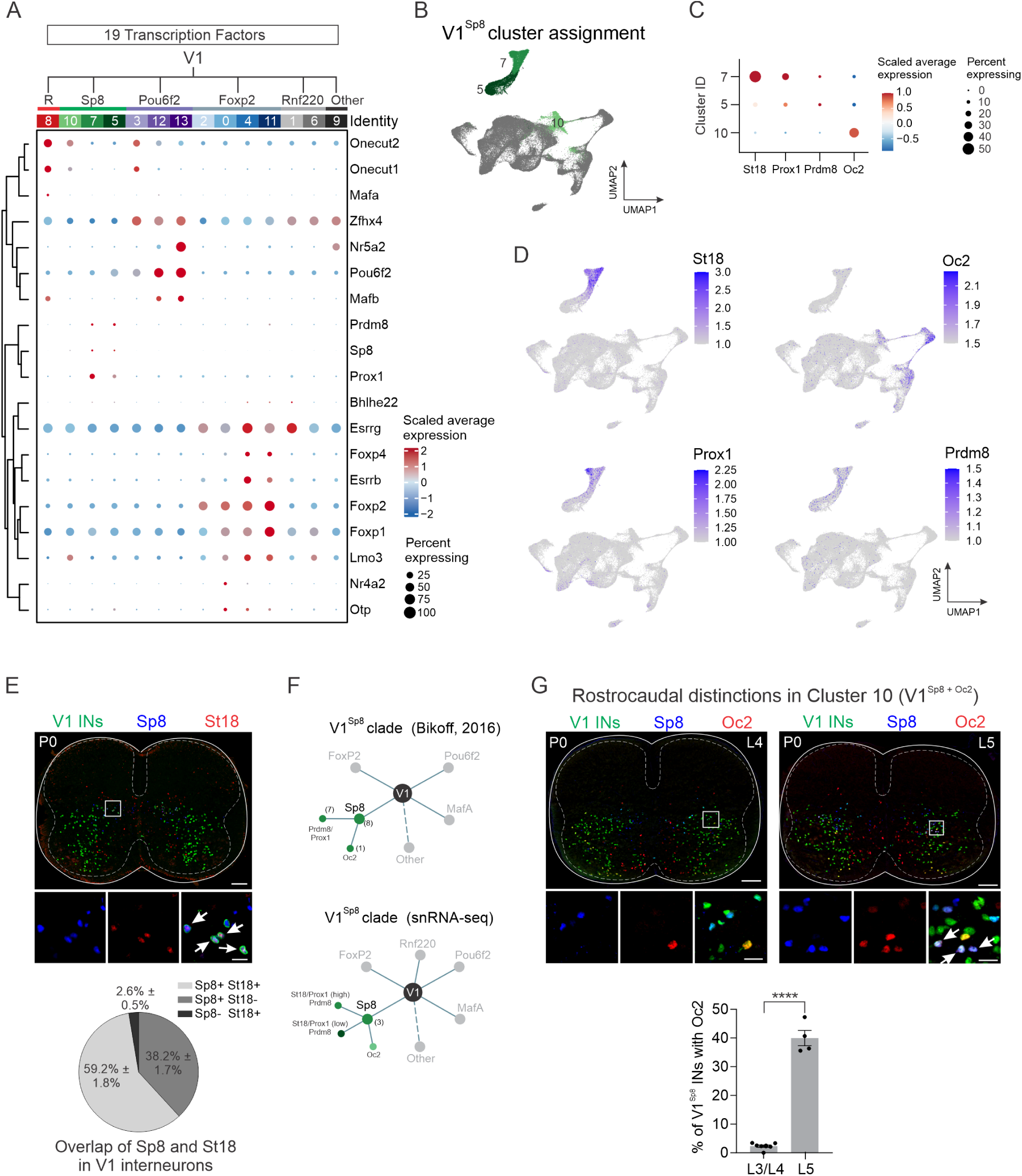
Analysis of transcription factor expression and V1^Sp8^ clade structure, related to Figures 1 and 3. (A) Dot plot showing scaled average expression of the 19 transcription factors (TFs) described previously in Bikoff et al.,^30^ all of which were validated via immunohistochemistry (IHC). Note that several genes were detected at low levels but still showed biased expression. (B) V1^Sp8^ cluster assignment, as described in Figure 1F. (C) Corresponding dot plot highlighting St18, Prox1, Prdm8, and Oc2 expression within the V1^Sp8^ clade. (D) UMAP plots of St18, Prox1, Prdm8, and Oc2 showing expression in V1 interneurons. (E) Top, IHC validation of St18 expression in V1 interneurons (arrows) in P0 lumbar spinal cord of *En1::Cre; RC.lsl.Sun1-sfGFP* mice. Scale bars = 100 μm (top) or 20 μm (inset). Bottom, the proportion of Sp8 and St18 co-expression in V1 interneurons as assessed by IHC. (F) Top, simplified V1^Sp8^ clade diagram adapted from Bikoff, et al.^30^ showing delineation of V1^Sp8^ neurons into seven Prdm8/Prox1 subsets and a single Oc2-expressing subset. Bottom, revised clade diagram based on snRNA-seq data, showing general alignment with previous analyses. (G) Top, IHC analysis of cluster #10 in P0 lumbar spinal cord of *En1::Cre; Tau.lsl.nLacZ* mice. Scale bars = 100 μm (top) or 20 μm (inset). Bottom, V1^Sp8+Oc2^ interneurons (arrows) were significantly enriched in caudal (L5) spinal segments compared to more rostral (L3/L4) segments, thus highlighting segmental differences in V1 cell types. \*\*\*\**p* < 0.0001, unpaired *t*-test.

**Figure S4.**
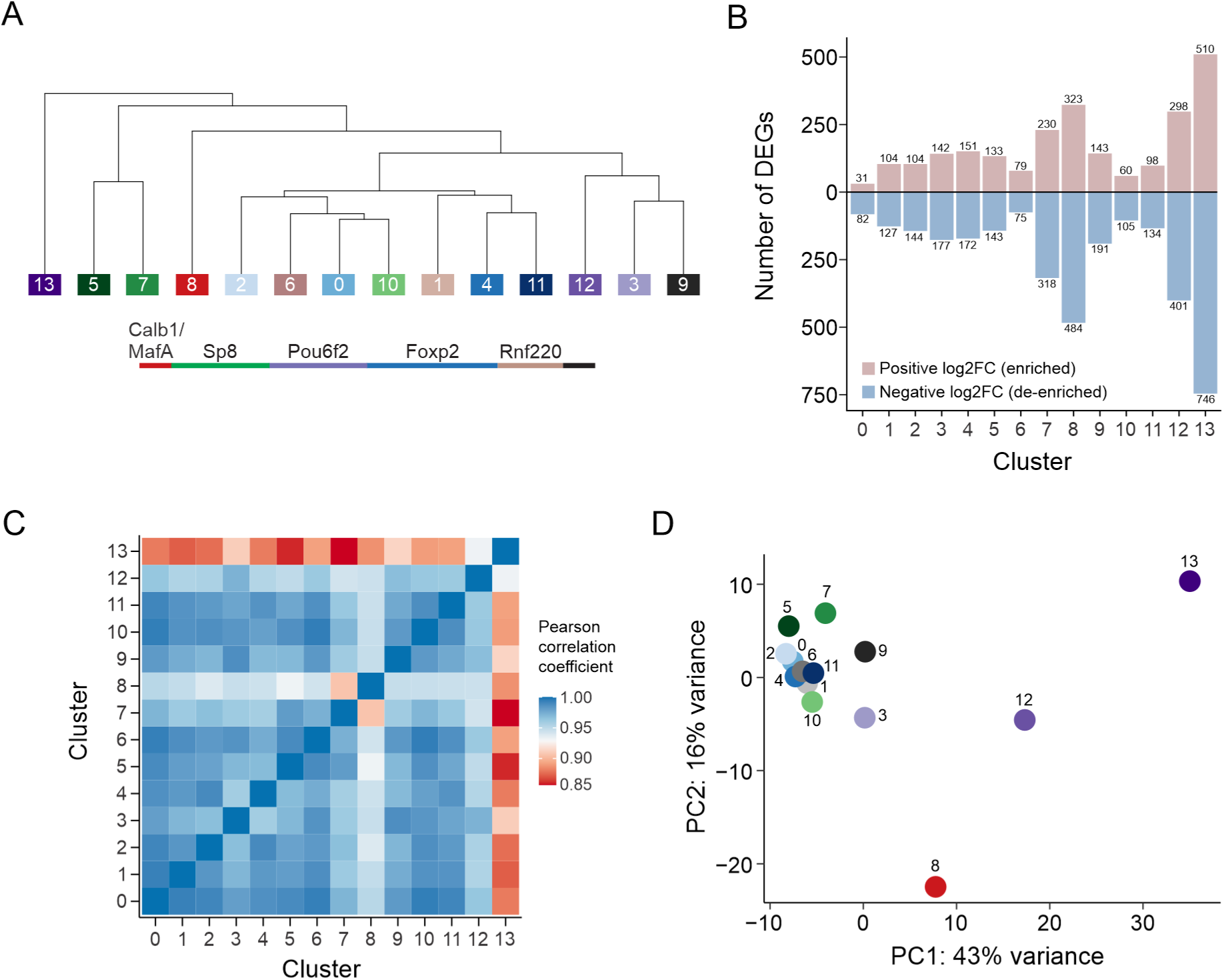
Taxonomic relationship and similarity of V1 clusters, related to Figure 3. (A) Dendrogram showing phylogenetic analysis based on the top 3000 most variable genes. (B) Number of differentially expressed genes (DEGs) per cluster based on non-parametric Wilcoxon rank sum test for differential gene expression. (C) Heatmap of Pearson correlation coefficients based on the log-normalized average expression of all genes per cluster identified cluster #13 as unique. (D) Principal component analysis (PCA) plot of aggregated counts per cluster showing that cluster #13 drives most of the variance along PC1.

**Figure S5.**
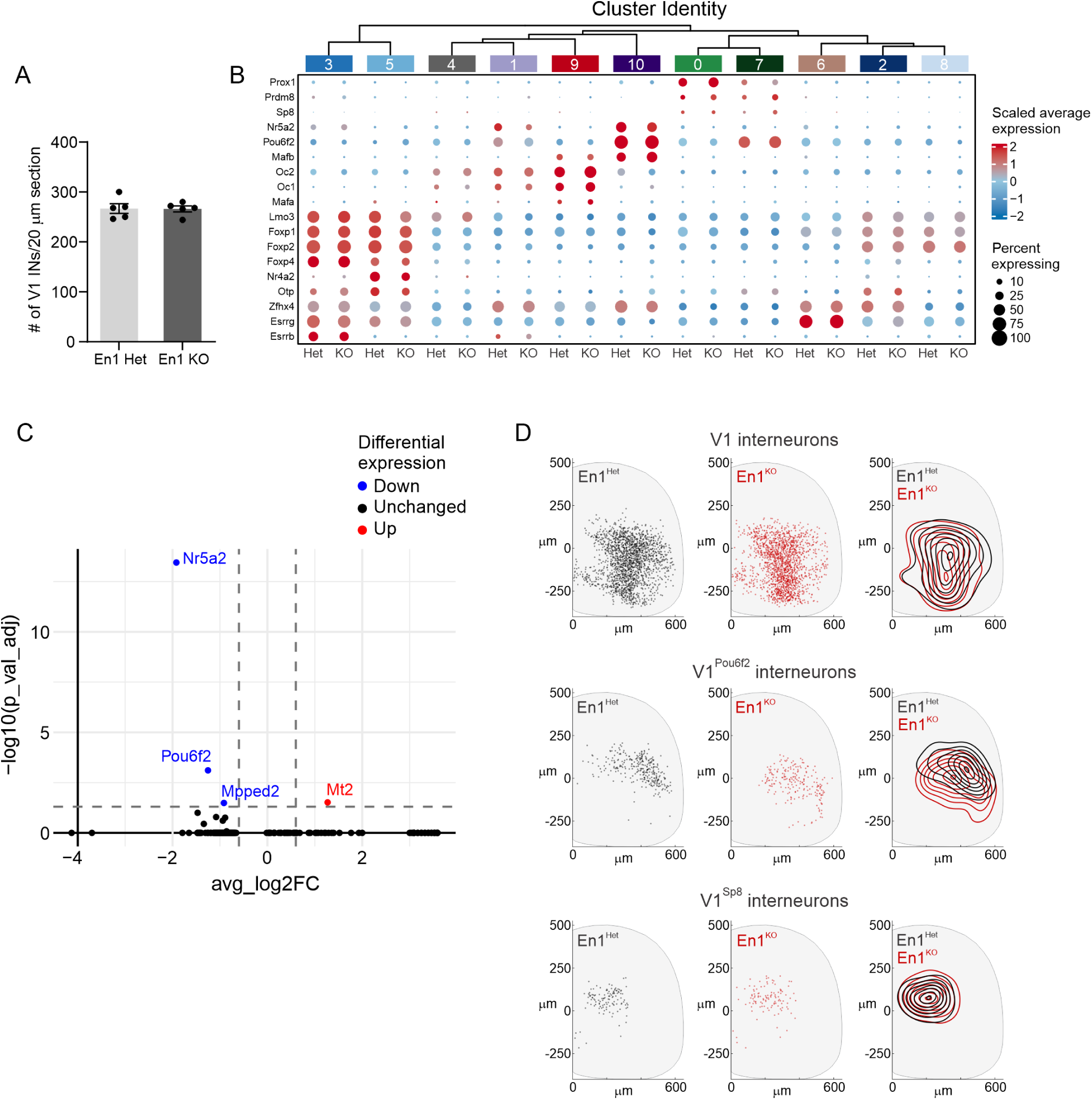
Additional gene expression analysis in *En1^Het^* and *En1^KO^* V1 interneurons, related to Figure 5. (A) The total number of V1 interneurons is similar in *En1^Het^* and *En1^KO^* mice (*p* = 0.815, unpaired two-tailed *t*-test). (B) Dot plot showing similar gene expression of 19 transcription factors in V1 nuclei from *En1^Het^* and *En1^KO^* mice. Note that the remaining *En1^KO^* V1 nuclei in cluster #10 retained a normal *Pou6f2^+^* and *Nr5a2^+^* identity. (C) Pseudo-bulk analysis of differentially expressed genes revealed that *Nr5a2* and *Pou6f2* are significantly downregulated in *En1^KO^* V1 nuclei, compared with that in *En1^Het^* nuclei. (D) Spatial distributions of the overall V1 interneuron population (top), V1^Pou6f2^ interneurons (middle), and V1^Sp8^ interneurons (bottom) in P0 lumbar spinal segments of *En1^Het^* (black) and *En1^KO^* (red) mice. The position of V1^Pou6f2^ neurons was altered in *En1^KO^* mice, whereas the position of V1^Sp8^ interneurons was unaffected (2-dimensional Kolmogorov-Smirnov test: *p* = 6.5 × 10^-13^ for V1^Pou6f2^ neurons*; p* = 0.095 for V1^Sp8^ neurons). Distributions were based on the following sample size: *En1^Het^* (V1): n = 4 animals, 13 hemisections, 1776 neurons from L3-L5 segments; *En1^KO^* (V1): n = 5 animals, 13 hemisections, 1702 neurons from L3-L5 segments; *En1^Het^* (V1^Pou6f2^): n = 8 animals, 21 hemisections, 309 neurons from L3 segments; *En1^KO^* (V1^Pou6f2^): n = 5 animals, 21 hemisections, 213 neurons from L3 segments; *En1^Het^* (V1^Sp8^): n = 3 animals, 8 hemisections, 122 neurons from L3-L5 segments; and *En1^KO^* (V1^Sp8^): n = 3 animals, 6 hemisections, 113 neurons from L3-L5 segments.

**Figure S6.**
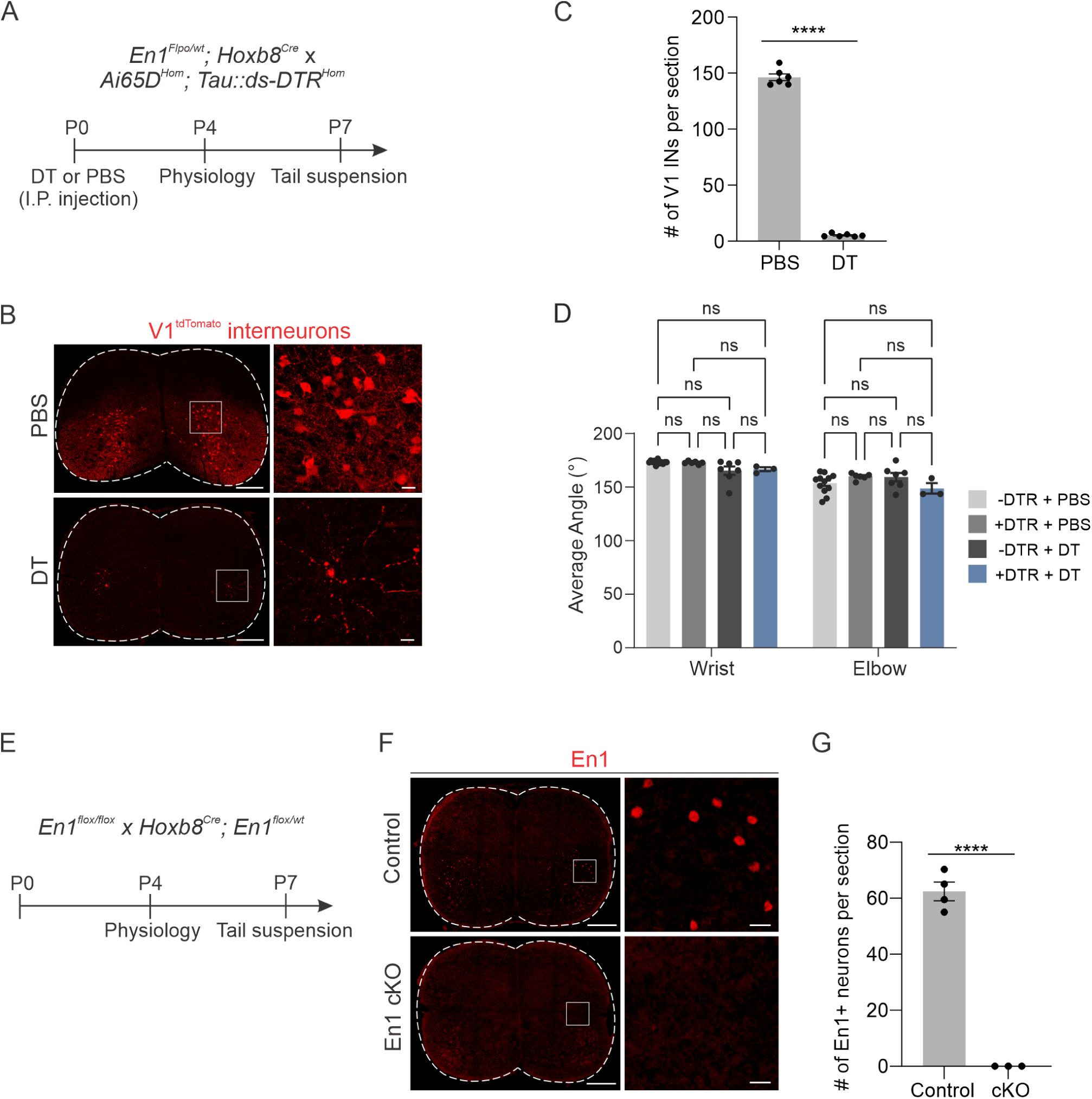
Validation of V1 ablation and *En1* conditional knockout experiments, and additional behavioral data, related to Figure 6. (A) Schematic of the V1 ablation paradigm. Diptheria toxin (DT) was administered at P0, followed by either physiological analysis via fictive locomotion 4 days post-DT administration, or limb kinematic analysis during tail suspension 7 days post-DT administration. All mice in the study were heterozygous for the *Tau::ds-DTR* and *Ai65D* alleles. Littermates lacking Cre, Flpo, or both were pooled and used as controls. (B-C) Immunohistochemical analysis and quantification in P7 lumbar spinal cords of control (+PBS) or V1-ablated (+DT) quadruple-heterozygous mice showed near complete ablation of V1 interneurons at P7 (n = 6 mice, \*\*\*\**p* < 0.0001, unpaired two-tailed *t*-test). Scale bars = 200 μm or 20 μm (inset) (D) The average angles of the wrist and elbow joints were not significantly changed upon ablation of V1 interneurons (n = 3-12 mice, mean ± SEM, two-way ANOVA followed by Tukey HSD test, ns = not significant). (E) Schematic of the experimental paradigm for physiological and behavioral analyses of *En1* conditional knockout (*En1 cKO*) mice. (F-G) Immunohistochemical analysis (F) and quantification (G) showing complete loss of En1 expression in *En1 cKO* (*Hoxb8::Cre; En1^flox/flox^*) mice in P0 lumbar spinal cord (n = 3-4 mice, \*\*\*\**p* < 0.0001, unpaired two-tailed *t*-test). Scale bars = 200 μm or 20 μm (inset).

## TABLES

**Table S1. Quality control metrics for CellRanger, related to Figures 1-5.**

**Table S2. Cell-type classification markers, related to Figures 1-5.**

**Table S3. Cell barcodes of analyzed V1 nuclei, related to Figures 1-5.**

**Table S4. Differentially expressed genes in V1 clusters, related to Figures 1-3.**

**Table S5. Differentially expressed genes by developmental age, related to Figure 4.**

**Table S6. Differentially expressed genes in V1 clusters in the *En1^Het^* versus *En1^KO^* experiment, related to Figure 5.**

## VIDEOS

Video S1. Examples of tail suspension assay in control, V1-ablated, and *En1 cKO* mice, related to Figure 6.

## EXPERIMENTAL METHODS

### RESOURCE AVAILABILITY

#### Lead contact

Further information and requests for resources and reagents should be directed to and will be fulfilled by the lead contact, Jay Bikoff.

#### Materials availability

The study did not generate new unique reagents.

#### Data and code availability

Raw sequencing data, counts tables, and associated metadata are deposited in the Gene Expression Omnibus (GEO) with accession number GEO: GSE275595. The code used for analysis of sequencing data is available at Github (https://github.com/Bikoff-Lab/V1_interneuron_snRNAseq_Age_Atlas and https://github.com/Bikoff-Lab/V1_KO_analysis).

### EXPERIMENTAL MODEL DETAILS

All experiments and procedures were performed in accordance with National Institutes of Health (NIH) guidelines and approved by the Institutional Animal Care and Use Committee (IACUC) of St. Jude Children’s Research Hospital. Mice were housed on ventilated racks in a temperature and humidity-controlled room on a standard 12-hour light/dark cycle with *ad libitum* food and water. Mice of both sexes were used for experiments. The following previously published mouse lines were used in this study: C57BL/6J (Jackson Laboratory #000664), *RC.lsl.tdT* (*Ai14*) (Jackson Laboratory #007914), *RC.dual.tdT* (*Ai65D*) (Jackson Laboratory #021875), *En1::Cre* (Jackson Laboratory #007916), *En1::Flpo,*^96^ *En1^flox^* (Jackson Laboratory #007918), *Hoxb8::Cre,*^75^ *Piezo2::Cre* (Jackson Laboratory #027719), *RC.lsl.Sun1-sfGFP* (INTACT) (Jackson Laboratory #021039), *Tau.lsl.nLacZ* (Jackson Laboratory #021162), and *Tau.ds.DTR.*^18^

### METHOD DETAILS

#### Tissue Collection and Nuclei Preparation

Spinal cord tissue was isolated from cervical, thoracic, and lumbar regions of *En1::Cre; RC.lsl.Sun1-sfGFP* double-heterozygous mice at P0, P14, P28, or P56 (for the developmental snRNA-seq atlas) or from the whole spinal cord (for P0 *En1^Het^* and *En1^KO^* experiments and the E13.5 reference sample), pooling 4-12 animals per experiment. Two independent biological replicates were performed per condition. Animals were anesthetized either on ice (P0 pups) or via intraperitoneal injection of Avertin (Tribromoethanol, 240 mg/kg body weight for P14, P28, and P56 animals). Transcardial perfusion was performed with ice cold, oxygenated artificial cerebral spinal fluid (ACSF) consisting of 125 mM NaCl (Sigma #S9888), 2.5 mM KCl (Sigma #9333), 1.25 mM NaH_2_PO_4_·H_2_O (Sigma #S71507), 26 mM NaHCO_3_ (Sigma #S5761), 25 mM glucose (Sigma #G7021), 5 mM ethyl pyruvate (Sigma #E47808), and 0.4 mM sodium ascorbate (Sigma #11140), supplemented after oxygenation with 1 mM MgCl_2_ (Sigma #M2392), 2 mM CaCl_2_ (Sigma #C5670), and blockers of neuronal activity and transcription (20 μM AP5 (Tocris #1060), 20 μM DNQX (Tocris #018910), 0.1 μM TTX (Tocris #1069), 5 μM Actinomycin D (Sigma #A1410)). Spinal cords were quickly extruded, separated into cervical, thoracic, and lumbar regions, then flash frozen and stored in liquid nitrogen until dissociation.

Nuclei dissociation was performed as described in Matson et al.^99^ Briefly, frozen tissue was mechanically dissociated with a Dounce homogenizer (Kontes Dounce Tissue Grinder) in 500 μL sucrose buffer (0.32 M sucrose, 10 mM HEPES pH 8.0 (Gibco #15630), 5 mM CaCl_2_ (Sigma #C5670), 3 mM magnesium acetate (Sigma #63052), 0.1 mM EDTA (Invitrogen #AM9261), 1 mM DTT with 0.1% Triton-X100 (Sigma #T8787), using five strokes of the A pestle followed by 15 strokes with the B pestle or until the tissue was dissociated. The resulting homogenate was filtered through a 40 μm strainer and washed with 3 mL of sucrose buffer. Nuclei were then centrifuged at 3200 × *g* for 10 minutes at 4°C. The supernatant was decanted, and nuclei were resuspended in 3 mL of sucrose buffer and transferred to an Oak Ridge centrifuge tube. A high-density sucrose solution containing 1M sucrose, 10 mM HEPES pH 8.0, 3 mM magnesium acetate, and 1 mM DTT was carefully layered underneath the remaining supernatant and then spun at 3200 x *g* for 20 minutes at 4°C. The supernatant was discarded, and the remaining nuclei were resuspended in 500 μL of phosphate-buffered saline (PBS) with 0.4 mg/mL BSA (NEB #30281) and 0.2U/μL RNAse inhibitor (Lucigen #30281-1) for subsequent processing by flow cytometry.

#### Flow Cytometry

Fluorescence-activated cell sorting (FACS) of sfGFP^+^ nuclei was performed using a FACSAria Fusion Cell Sorter (BD Biosciences, San Jose, CA). The FACSAria Fusion is equipped with a 50 mW 488nm laser and a 530/30 bandpass filter and uses BD FACSDiva software v8.0.1. GFP^+^ nuclei were selected based on log side scatter (SSC-A) versus log GFP fluorescence (GFP-A) and sorted in purity mode (sheath pressure 21 PSI, 100 um nozzle) into 1.5 mL Protein LoBind Eppendorf tubes coated overnight with PBS +2% BSA. Samples were visually inspected and counted on a hemocytometer after processing by flow cytometry.

#### snRNA-seq Library Preparation and Sequencing

Sorted GFP^+^ nuclei were loaded onto the 10x Genomics Chromium Controller (#1000171) to generate single-nucleus gel beads-in-emulsions (GEMs) using the Chromium Next GEM Single Cell 3’ GEM, Library, and Gel Bead Kit v3.1 (#1000121) and Chromium Single Cell B Chip Kit (#1000120), according to the manufacturer’s instructions. Approximately 8,000 nuclei were loaded for capture per sample. Following capture and lysis, cDNA synthesis, amplification, and library construction were performed on an ABI ProFlex PCR System (Applied Biosystems). The libraries were quantified using the Quant-iT Pico Green dsDNA assay (Thermo Fisher) and through low-pass sequencing with a Miseq nano kit (Illumina). Quality assessments were conducted using the 4200 TapeStation D1000 and D5000 ScreenTape assay (Agilent). Paired-end 100-cycle sequencing (Run configuration: 101-8-8-101) was carried out on a NovaSeq 6000 (Illumina) to a depth of at least 30,000 reads per nucleus in the St. Jude Hartwell Center for Biotechnology.

#### snRNA-seq Analysis

##### Data preprocessing and quality control

Sequencing data were aligned to the mouse genome (mm10, 10x Genomics version 2020 A) using CellRanger (v7.0.0) with default parameters except for the --force-cells option, which was set to 8,000 or 9,000 for all datasets based on the estimated number of nuclei loaded by manual counting after sorting. The E13.5 dataset used for positive selection was mapped using CellRanger v3.0.0. Ambient RNA was removed from the samples using SoupX^37^ (version 1.6.2). Of the 24 samples used for the atlas, one (P56LR1) was removed at this stage due to high ambient RNA identified by SoupX. The gene expression matrix was then analyzed in R (v4.1.2) using the Seurat package (v4.3.0). See https://github.com/Bikoff-Lab for the complete list of libraries and version numbers used for downstream data processing. Each dataset underwent individual quality control (QC) to remove low-quality data and doublets. Briefly, nuclei were filtered out if they contained <1000 unique molecular identifiers (UMIs), >2.5% of transcripts derived from mitochondrial genes, or were above or below ±3 median absolute deviations (MADs) from the median number of UMIs and genes per sample. Any gene that appeared in <10 cells was removed.

##### Identification of V1 nuclei

To generate the V1 interneuron atlas, nuclei that passed QC underwent an iterative process of negative and positive selection to remove any contaminating non-V1 nuclei. Initial rounds of negative selection were performed first on a merged Seurat object, followed by negative selection on an integrated dataset with batch correction. We used a combination of QC metrics and known marker genes to remove non-V1 nuclei. Positive selection was performed using a previously annotated dataset generated by Delile et al.^22^ and the label transfer method in Seurat. A brief summary of this process is outlined below.

###### Negative selection

Initial rounds of negative selection on a simple merge of all samples first underwent standard log normalization per 10,000 reads followed by scaling of all genes. The top 2000 genes with the most variable expression across the entire merged dataset using the vst method and then linear dimensionality reduction with PCA was performed. To determine how many principal components to use in downstream analysis, two criteria were used: (1) the cumulative percent variation explained was >90% and the individual percent variation explained was <5%; (2) The change in percent variation was >0.1% (custom function called find_PC). Further dimensionality reduction and clustering were performed in accordance with the standard Seurat workflow (FindNeighbors, FindClusters, and RunUMAP) using default parameters. To annotate the resulting clusters and eliminate non-V1 nuclei, unhealthy/dying cells and empty GEMs were identified by using a combination of the total number of counts, total number of unique genes detected, percent mitochondrial reads, and the number of DEGs. A series of marker genes was used to identify non-neuronal clusters and defined types of neurons, based on previous studies.^22,25,28,39,81^ (See Table S1 for a list of marker genes used to identify these cell types). After removing non-V1 nuclei, the negative selection process was repeated three times until no further contamination was identified. The resulting data were subjected to further negative selection by using a similar approach but with batch correction using Seurat’s anchor-based integration method. Standard normalization and scaling of the counts was followed by identification of the top 2000 most variable genes for each dataset, which were then cross-referenced across all samples to find common variable features across datasets. This information was used to perform Seurat’s FindIntegrationAnchors and IntegrateData to create a batch-correct dataset that then underwent standard dimensionality reduction and clustering as described above. Annotation of any remaining non-V1 nuclei was performed using the same marker genes outlined above. These nuclei were removed from the dataset, and this process was repeated until no further contamination was identified.

###### Positive selection

The remaining nuclei that passed QC and multiple rounds of negative selection then underwent positive selection using Delile et al. as a reference. Starting with fastq files, this reference dataset was re-analyzed using the above workflow, with cell types manually identified using the marker genes described in the original publication. A single embryonic snRNA-seq experiment was performed to obtain an age-matched E13.5 dataset enriched for V1 nuclei. After QC of this dataset, the Seurat workflow was used to annotate the query dataset with labels from the reference dataset. The E13.5 snRNA-seq dataset was then used for positive selection of the postnatal V1 interneuron nuclei already filtered by QC metrics and negative selection markers using the FindTransferAnchors method in Seurat. Nearly all nuclei (89494 of 89545) that passed negative selection were called as V1 neurons by this label-transfer method.

Barcodes of the identified V1 nuclei were subset from the original count matrix and re-analyzed via SCT normalization and integrated batch correction. Briefly, the raw counts were log normalized and scaled per usual. SCTransform was performed for normalization and variance stabilization on each dataset individually by using the “glmGamPoi” method. The SCT-normalized object was then integrated using the standard Seurat workflow, using the P0 dataset as a reference to reduce computation time. The integrated object then underwent standard dimensionality reduction and clustering, as described above, using one of two resolutions: 0.003 to produce two clusters (Figure 1B) or 0.24 to produce 14 clusters (Figures 1-4).

For P0 whole spinal cord samples collected from *En1^Het^* and *En1^KO^* animals that formed the basis of Figure 5, two independent biological replicates were independently processed and analyzed in Python using Scanpy (v1.9.8). Briefly, after filtering low-quality nuclei, doublets, and rarely detected genes using the QC cutoffs described above, the data was log-normalized and scaled with batch differences regressed out. No further integration was needed. Non-V1 clusters of nuclei were identified and removed using the marker genes listed in Supplemental Table 1. This dataset did not undergo positive selection.

##### Assignment of clade identity

To assign a particular cluster to a V1 clade, the expression of four clade markers (*Foxp2, Sp8, Pou6f2*, or *Chrna2* as a proxy for *MafA*) was assessed at the single-nucleus level and on a per-cluster basis. At the single-nucleus level, a nucleus was considered positive if its log-normalized expression was > 2.8 for *Foxp2*, >0.8 for *Sp8*, >2.6 for *Pou6f2*, or >1.1 for *Chrna2*. The average gene expression was then calculated across all nuclei within each cluster, the average gene expression was scaled, and z-scores were used to assign cluster identity. Positive z-scores were typically sufficient to assign a cluster to a single clade. Only cluster #5 had two positive z-scores, for which the higher z-score (corresponding to Sp8) was used to determine clade identity.

##### Differential gene expression to find cluster-specific genes

Differential expression was performed in Seurat with either ‘FindAllMarkers’ or ‘FindMarkers’ using a Wilcoxon sum rank test and default parameters. All tests were done using the Wilcoxon rank sum test. *P* values were adjusted using the Bonferroni method based on the number of genes in the dataset. Top markers were ranked by their log fold change (FC), with log_2_FCs > ±0.25 considered significant.

##### Analysis of cluster relationships

The relationships between V1 nuclei in different clusters were assessed using several methods. To generate dendrograms, an agglomerative approach was used to examine hierarchical relationships among clusters via the BuildClusterTree algorithm in Seurat on the integrated, scaled data. To evaluate the relationships among clusters based on overall gene expression, Pearson correlation coefficients were calculated on the average expression of all log-normalized genes within each cluster using the cor function in R. Relationships among clusters in PC space were assessed by aggregating counts for all clusters and running them through the DEseq2 bulk RNAseq pipeline.^98^ Local inverse Simpson’s Index (LISI) scores were used to assess intermixing of nuclei across developmental age within clusters, calculated with the compute_Lisi function using two-dimensional UMAP coordinates for input.^100^

##### Gene Ontology analysis

Age-related changes in gene expression were identified using the default Wilcoxon Rank Sum test in Seurat for nuclei isolated from P0 animals versus all other ages, and for P28 and P56-derived nuclei versus all other time points due to their similarity in gene expression. Genes that were positively enriched with a corrected *p* value <0.05 were imported into ShinyGO (v0.80) for identification of the most significant gene ontology (GO) terms for Biological Processes.^101^

##### Compositional analysis of nuclei in clusters

<u>S</u>ingle-<u>c</u>ell <u>c</u>ompositional <u>d</u>ata <u>a</u>nalysis (scCODA) was used to determine significant changes in nuclei composition between *En1^Het^* and *En1^KO^* snRNA-seq samples.^102^ scCODA uses Bayesian modeling based on a hierarchical Dirichlet-Multinomial distribution to accommodate the limitations of snRNA-seq experiments on compositional analysis, including limited sample size, biases in cell-type correlation estimates, and the sparsity of cell-type specific manipulations. Cluster #2 was selected as a reference because it was relatively abundant in the dataset, it exhibited the smallest absolute change in composition across all four samples, and it had the least dispersion of relative abundance over all samples. Using a FDR of 0.05, only cluster #10 had a credible effect on cell type composition with a log_2_FC of −1.61 and an inclusion probability parameter of 0.97.

##### Pseudo-bulk analysis

Pseudobulk analysis was performed via summation of raw counts from all nuclei within a sample of interest. Aggregated counts were then analyzed similar to bulk RNAseq data using the DEseq2 pipeline. Briefly, a DESeq2 object was created, QC was assessed by PCA and hierarchical clustering, and then count normalization was performed and differential gene expression was analyzed using the Wald test.

#### Immunohistochemistry, RNAscope In Situ Hybridization, and Confocal Imaging

Mice were deeply anesthetized via intraperitoneal injection of Avertin (Tribromoethanol, 240 mg/kg body weight) or on ice (for early postnatal animals) and transcardially perfused with PBS followed by ice-cold 4% paraformaldehyde (EMS #15714, diluted in PBS). Spinal cords were removed and post-fixed for 2 hours (neonatal spinal cords) or overnight (P14, P28, and P56 spinal cords) at 4°C, followed by cryoprotection in 30% sucrose (w/v) in 0.1M PB for at least 24 hours at 4°C. Tissue was embedded in OCT (Fisher #23730571), frozen, cryosectioned in the transverse plane at 20 μm, and mounted on Superfrost Plus slides (Fisher #1367811E). IHC was performed through sequential exposure to primary antibodies overnight at 4°C, and fluorophore-conjugated (Alexa Fluor Plus 488 or DyLight 488, Alexa Fluor Plus 555 or Cy3, and Alexa Fluor Plus 647 or Cy5) secondary antibodies for 1 hour at room temperature. Sections were mounted using Fluormount-G (SouthernBiotech) or ProLong Diamond (Invitrogen P36970 without DAPI or P36971 with DAPI) and coverslipped for imaging. Confocal images were acquired on a Lecia SP8 AOBS (Leica Microsystems) confocal microscope using a 20×/0.8 NA objective, or a LSM 710 Meta Confocal microscope (Carl Zeiss) using a Plan-Apochromat 20×/0.8 M27 objective.

The following primary antibodies were used: chicken anti-β-Galactosidase (1:5000, Abcam #ab9361); rabbit anti-Calbindin D28K (1:2000, Swant #CB38); guinea pig anti-dsRed (1:16000, Jessell Lab); guinea pig anti-En1 (1:16000, Jessell Lab); guinea pig anti-FoxP1 (1:20000, Jessell Lab); goat anti-Foxp2 (1:30000, Abcam #ab1307); goat anti-FoxP2 (1:500, Santa Cruz #sc-21069); chicken anti-GFP (1:30000, Aves #GFP-1020); rabbit anti-GFP (1:1000, Invitrogen #A-11122); guinea pig anti-Lmo3 (1:8000, Jessell Lab); rabbit anti-MafA (1:2000, Novus #NB400-137); rabbit anti-MafB (1:2000, Sigma #HPA005653); mouse anti-Nr3b3 (1:2000, R&D Systems #PP-H6812-00); rabbit anti-Nr4a2 (1:500, Santa Cruz #sc-5568); goat anti-Nr5a2 (1:100, Santa Cruz #sc-21132); rabbit anti-Oc1 (1:200, Sigma #HPA003457); sheep anti-Oc2 (1:2000, R&D Systems #AF6294); guinea pig anti-Otp (1:16000, Jessell Lab); rat anti-Pou6f2 (1:16000, Jessell Lab); guinea pig anti-Prdm8 (1:10000, Jessell Lab); rabbit anti-Prox1 (1:2000, Millipore #AB5475); rabbit anti-Rnf220 (1:5000, Novus #NBP1-88487); goat anti-Sp8 (1:2000, Santa Cruz #sc-104661); and rabbit anti-Zfhx4 (1:1000, Sigma #HPA023837). Antibodies generously provided as gifts include rabbit anti-Nr3b2 (1:2000, from Jeremy Nathans) and rat anti-St18^40^ (1:8000, from Guoqiang Gu).

RNAscope ISH was performed according to the manufacturer’s instructions (Advanced Cell Diagnostics, CA) with the following modifications: drying time, 1 hour; antigen retrieval, 5 minutes; protease treatment, 20 minutes. The following probes were used: Bcl11a (#563701), Dach1 (#412071), Kcnq5 (#511131), Kcnh7 (#1007281), Pax8 (#574431), St18 (#443271), and tdTomato (#317041). Signal was developed using Opal fluorophores (Akoya Biosciences) at a 1:1000 dilution, and slides were mounted in ProLong Gold (Invitrogen #P36930). Fluorescent images were acquired on a Leica TCS SP8 STED confocal microscope in resonant mode using a 63×1.4 oil objective with a 2× zoom.

#### Analysis of Interneuron Spatial Distributions

The position of V1 interneurons in P0 lumbar spinal segments of *En1::Cre* heterozygous; *Tau.lsl.nLacZ* (*En1^Het^*) or *En1::Cre* homozygous; *Tau.lsl.nLacZ (En1^KO^*) mice was analyzed as previously described, using the “Spots” function in Imaris (Bitplane, Oxford Instruments Andor).^30^ Briefly, Cartesian coordinates for each interneuron were determined in the transverse spinal cord plane, with respect to the midpoint of the central canal, defined as position (0,0). All sections were normalized to a standardized spinal cord hemisection to account for variations in spinal cord dimensions along the rostrocaudal axis (distance from the central canal to the lateral boundary: 650 μm; distance from central canal to bottom-most boundary: 400 μm). Coordinates were plotted using custom Matlab scripts, with contours constructed using the kde2d function. Differences in spatial distributions were assessed using the two-dimensional Kolmogorov-Smirnov test implemented with the Python function ndtest (https://github.com/syrte/ndtest).

#### Fictive Locomotion

Fictive locomotion experiments were performed on P0 to P4 mice of both sexes. Briefly, after decapitation, mice were eviscerated and then transferred to a dissecting chamber containing cold aCSF consisting of 4 mM KCl, 128 mM NaCl, 21 mM NaHCO_3_, 0.5 mM NaH_2_PO_4_ ● H_2_O, 30 mM glucose, 1 mM CaCl_2_, and 1 mM MgSO_4_ and oxygenated with 95% O_2_/5% CO_2_. After a dorsal laminectomy, the spinal cord and attached roots were continuously perfused with room temperature oxygenated aCSF containing 4 mM KCl, 128 mM NaCl, 21 mM NaHCO_3_, 0.5 mM NaH_2_PO_4_ ● H_2_O, 30 mM glucose, 2 mM CaCl_2_, 1 mM MgSO_4_, 10 µM 5-HT (Sigma #H7752), and 5 µM NMDA (Sigma #M3262). The spinal cords were allowed to recover for 30 minutes at room temperature before recording. Extracellular recordings from ventral roots were obtained using tight-fitting glass suction electrodes (for smaller roots Drummond #3-000-203-G/X, for larger roots A-M Systems #592800) from either the flexor-dominated left or right L2 root and/or the extensor-dominated left or right L5 root. Signals were amplified 10,000 times, bandpass filtered between 0.1 Hz and 5000 Hz, digitized at 10 kHz (Digidata 1550B; Molecular Devices), and collected using pClamp software. To calculate fictive locomotor frequency, traces were low-pass filtered using a moving mean of 10000 points (equal to 1 second of data) and the power spectral density was calculated using the periodogram function in Scipy. The frequency corresponding to the maximum power was used to determine locomotor speed.

#### Tail suspension and kinematic analysis

For tail suspension experiments, P7 mice were hung by there tails at an ambient temperature of 30°C and recorded facing their ventral side on a Basler acA720 camera at a rate of 200 frames/second for 6 minutes. Videos of tail-suspended markerless pups were tracked using DeepLabCut (DLC).^103^ A DLC network was trained to track nine landmarks on the pups: forepaw, wrist, elbow, armpit, hindpaw, ankle, knee, groin, and anus. Videos were then manually annotated to select frames in which all the landmarks were visible and the pups were not executing turning or flipping body movements. The DLC-tracked coordinates in these frames were then analyzed using AutoGaitA^104^ to assess the following joint angles across experimental groups: wrist angle (forepaw and elbow coordinates), elbow angle (wrist and armpit coordinates), ankle angle (hindpaw and knee coordinates), and knee angle (ankle and groin coordinates). All statistical analyses were performed using GraphPad Prism.

### QUANTIFICATION AND STATISTICAL ANALYSIS

Two-tailed *t*-tests (unpaired) were used to assess significance between means, with a Holm-Sidak correction for multiple comparisons where appropriate, as indicated in the figure legends. Two-way ANOVA followed by a Holm-Sidak test for multiple comparisons was used to assess significance between mean frequency of rhythmic activity and joint angle measurements upon V1 ablation, grouped by genotype and treatment condition. Differences among groups were considered significant if *p* < 0.05. *p* values are denoted by asterisks: \**p* < 0.05; \*\**p* < 0.01; \*\*\**p* < 0.001; \*\*\*\**p* < 0.0001; ns – not significant. Statistical analyses were performed using GraphPad Prism software and R, and all data are represented as mean ± SEM unless otherwise indicated.

